# Variation in maize chlorophyll biosynthesis alters plant architecture

**DOI:** 10.1101/2020.01.24.917732

**Authors:** Rajdeep S. Khangura, Gurmukh S. Johal, Brian P. Dilkes

**Affiliations:** Department of Botany and Plant Pathology, Purdue University, West Lafayette, Indiana 47907; Center for Plant Biology, Purdue University, West Lafayette, Indiana 47907; Department of Biochemistry, Purdue University, West Lafayette, Indiana 47907

**Author notes:** Corresponding Authors Brian P. Dilkes, 170 S University Ave, Purdue University, West Lafayette, Indiana 47907, Gurmukh S. Johal, 915 W State St, Purdue University, West Lafayette, Indiana 47907.

**Keywords:** protoporphyrin IX, tassel architecture, axillary meristem, sugar signaling, shade avoidance, metabolism

## Abstract

Chlorophyll is a tetrapyrrole metabolite essential for photosynthesis in plants. The *oil yellow1* (*oy1*) gene of maize encodes subunit I of Magnesium chelatase, the enzyme catalyzing the first committed step of chlorophyll biosynthesis. A range of chlorophyll contents and net CO_2_ assimilation rates can be achieved in maize by combining a semi-dominant mutant allele, *Oy1-N1989,* and cis-regulatory alleles encoded by the Mo17 inbred called *very oil yellow1* (*vey1*). We previously demonstrated that these allelic interactions can delay reproductive maturity. In this study, we demonstrate that multiple gross morphological traits respond to a reduction in chlorophyll. We found that stalk width, number of lateral branches (tillers), and branching of the inflorescence decline with a decrease in chlorophyll level. Chlorophyll variation suppressed tillering in multiple maize mutants including *teosinte branched1*, *grassy tiller1*, and *Tillering1* as well as the *tiller number1* QTL responsible for tillering in many sweet corn varieties. In contrast to these traits, plant height showed a non-linear response to chlorophyll levels. Weak suppression of *Oy1-N1989* by *vey1^B73^* resulted in a significant increase in mutant plant height. This was true in multiple mapping populations, isogenic inbreds, and hybrid backgrounds. Enhancement of the *Oy1-N1989* mutants by the *vey1^Mo17^* allele reduced chlorophyll contents and plant height in mapping populations and isogenic inbred background. We demonstrate that the effects of reduced chlorophyll content on plant growth and development are complex and that the genetic relationship depends on the trait. We propose that growth control for branching and architecture are downstream of energy balance sensing.

## Introduction

Land plants project their aerial parts into three-dimensional space to capture solar radiation and convert it into chemical energy via photosynthesis (Rabinowitch and Govindjee 1965; Govindjee *et al*. 2017). Energy is stored as carbon-carbon bonds and primarily transported as sugars from photosynthetically active tissues, such as leaves, to rapidly growing parts of the plant. The addition of biomass to young tissue is achieved by the mobilization of this photosynthetically fixed carbon (Yu *et al*. 2015). Thus, growth of the plant body is achieved by consuming excess sugars from photosynthesis. To sustain maximum growth rates, plants must coordinate growth and metabolism to prevent undershooting or overshooting the available resources (Lemoine *et al*. 2013). Unsurprisingly, energy deficits from manipulation of photosynthesis can result in dramatically decreased growth.

One way to manipulate photosynthetic activity and energy resources in plants is by altering chlorophyll biosynthesis (Walker *et al*. 2017; Khangura *et al*. 2019a). Magnesium chelatase (MgChl) is a hetero-oligomeric enzyme complex that catalyzes the first committed step of chlorophyll biosynthesis by conversion of protoporphyrin IX (PPIX) into magnesium-PPIX (Walker and Weinstein 1991; Gibson et al. 1995). In maize, one of the subunits of the MgChl complex is encoded by the *oil yellow1* gene (*oy1*; B73 RefGenV3: GRMZM2G419806; B73 RefGenV4: Zm00001d023536). This gene received its name because weak hypomorphic alleles result in a light green to yellow color reminiscent of vegetable oil (Eyster 1933; Sawers *et al*. 2006). We previously demonstrated that the semi-dominant *Oy1-N1989* mutant was strongly modified by a QTL called *very oil yellow1* (*vey1*) that is likely encoded by cis-acting regulatory polymorphisms at *oy1* itself (Khangura *et al*. 2019a). The highly heritable *vey1* QTL is a major modifier of chlorophyll contents in heterozygous *Oy1-N1989/+* mutants but had no effect on chlorophyll accumulation in wildtype plants from either B73 x Mo17 biparental mapping populations or a maize association panel (Khangura *et al*. 2019a). The *vey1* alleles are detectable as a cis-acting eQTL and generate allele-specific expression at *oy1* (Li *et al*. 2013, 2018; Waters *et al*. 2017; Khangura *et al*. 2019a, 2019b). These expression level differences at *oy1* are visible at the mRNA level in both mutants and wildtypes, but the impact on chlorophyll content was only visible in the presence of the *Oy1-N1989* allele (Khangura *et al*. 2019a). This allelic interaction indicates the value of using mutants to uncover hidden genetic variation in maize. In the B73 x Mo17 populations, the *vey1* allele from B73 (*vey1^B73^*) was associated with higher expression of OY1, whereas the *vey1* allele from Mo17 (*vey1^Mo17^*) was linked with lower expression of OY1.

Using the *Oy1-N1989* mutant allele, and the two *vey1* modifier alleles from B73 and Mo17, we previously developed populations of maize to investigate the effects of this cryptic variant on chlorophyll content (Khangura *et al*. 2019a), net CO_2_ assimilation, and reproductive maturity (Khangura *et al*. 2019b). These three phenotypes showed high linear correlations with each other and were consistent with expression of the wildtype *oy1* allele such that greater expression of the wildtype OY1 transcript resulted in suppression of the *Oy1-N1989/+* mutant phenotype (Khangura *et al*. 2019a, 2019b). Our results, thus far, are consistent with an integrative measure of carbon assimilation linking energy status and vegetative to reproductive transition in maize (Minow *et al*. 2018).

In this report, we show that these allelic combinations at *vey1* and *oy1* can be used to study the effect of chlorophyll content and photosynthetic rate on plant growth and development. We again used crosses of *Oy1-N1989/+* mutants to B73 x Mo17 recombinant inbred lines (RIL) and near-isogenic lines (NIL) carrying alternate alleles at *vey1* to vary chlorophyll content in maize. QTL mapping was used to detect loci that modified stalk width and plant height in both wildtype and mutant F_1_ hybrid (RIL x tester) populations. The single locus testing of *oy1-vey1* interactions for these traits were tested directly using NIL x tester crosses. We found stalk widths of *Oy1-N1989* mutants were narrower than wildtype siblings and this was enhanced by the *vey1^Mo17^* allele, just as we previously found for chlorophyll content, net CO_2_ assimilation, and time to reproductive maturity (Khangura *et al*. 2019a, 2019b). To our surprise, we found that plant height in the *Oy1-N1989* mutants can be either greater or less than their congenic wildtype siblings. This divergence in plant height of heterozygous *Oy1-N1989* mutants was contingent upon the *vey1* allele. We also tested the effects of *Oy1-N1989* and *vey1* interaction on tassel phenotypes and other axillary meristem structures such as tiller numbers in isogenic materials. A previous study identified effects of a recessive partial loss-of-function allele at *oy1* on tassel branch number (Brewbaker 2015). Our results agree with Brewbaker (2015) as *Oy1-N1989* decreased tassel branch number dramatically in the presence of *vey1^Mo17^* allele and the effect was suppressed by *vey1^B73^*. Remarkably in high-tillering sweet corn, *Tillering1 (Tlr1)*, *teosinte branched1 (tb1)* and *grassy tillers1 (gt1)* mutants, *Oy1-N1989* reduced the number of tillers at maturity and an epistatic effect of *Oy1-N1989* on *tb1* and *gt1* was further enhanced by the *vey1^Mo17^* allele. The effect of a chlorophyll biosynthetic mutant on plant architecture and the unexpected opposing effect of the mutant and the *vey1* modifier on plant height are discussed.

## Materials and methods

### Plant materials

The *Oy1-N1989* mutant allele was obtained from the Maize Genetics Cooperation Stock Center (MGSC; University of Illinois, Urbana/Champaign, IL). The mutant heterozygotes (*Oy1-N1989*/*+*) were backcrossed as a pollen-parent for a minimum of seven times into B73 genetic background (*Oy1-N1989*/*+*:B73). For linkage mapping, *Oy1-N1989*/*+*:B73 pollen was crossed to 216 Intermated B73 x Mo17 recombinant inbred lines (IBM; Lee *et al*, 2002) and 251 Syn10 double haploid lines (Syn10; Hussain *et al*, 2007). A similar crossing scheme was used to develop F_1_ hybrid progenies for QTL validation with 35 B73-Mo17 near-isogenic lines (NIL; Eichten et al., 2011), consisting of 22 NIL with Mo17 donor introgressions into recurrent B73 (B73-like NIL) and 13 lines with B73 donor introgressions into recurrent Mo17 (Mo17-like NIL) genetic background. Each F_1_ hybrid progeny from these crosses segregated 1:1 for isogenic mutant (*Oy1*-*N1989*/*+*) and wildtype (*+*/*+*) siblings.

To test the effect of *Oy1-N1989* and its modifier *vey1* on tillering, we collected mutant stocks in isogenic B73 background. The genetic stock 127G harboring semi-dominant *Tillered1-N1590* allele (*Tlr1*) mutation (likely an additional allele of *tb1*; (Springer and Bennetzen 1995)) was obtained from MGSC in the Hi27 background. Stocks with the *tb1-ref* and *gt1-1* alleles (hereinafter *tb1* and *gt1*) in the B73 background were obtained from Dr. Clint Whipple (Brigham Young University, Provo, Utah, USA). To develop stocks segregating for both *vey1* and *tb1*, we first crossed a B73-like NIL (Eichten *et al*. 2011) homozygous for the *vey1^Mo17^* locus (Khangura, Marla, *et al*. 2019) to *tb1* mutants to generate *tb1*/+;*vey1^Mo17^*/*+* F_1_ progeny. Second, pollen from homozygous *tb1* mutants was used to pollinate *Oy1-N1989*/*+*:B73 ears to generate *tb1*/+; *Oy1-N1989*/+ F_1_ progenies. Third, the *tb1*/+; *Oy1-N1989*/+ plants from the first cross were used as pollen-parents in crosses to *tb1*/+; *vey1^Mo17^*/+ ears. The progeny of this cross segregated for all possible genotypes at the *tb1* and *oy1* locus. A similar crossing strategy was used for testing the effect of *Oy1-N1989* and *vey1* alleles on tillering in the *gt1* and *Tlr1* mutants.

### Location, Experimental design, and planting

All the field experiments were performed at the Purdue Agronomy Center for Research and Education (ACRE) in West Lafayette, Indiana. The test cross populations of IBM F_1_ and Syn10 F_1_ hybrid progenies were evaluated in 2013 (IBM) and 2014 (Syn10) in a randomized complete block design (RCBD) with one and two replications, respectively. The F_1_ progeny of *Oy1*-*N1989*/*+*:B73 crossed with B73 and Mo17 were used as parental checks in each range treated as a block. The F_1_ progenies of *Oy1*-*N1989*/*+*:B73 with NIL and parents (B73 and Mo17) were planted in a RCBD with five replications in 2016. All F_1_ families were planted as a single plot of 12-16 plants. Plots were sown in a 3.84 m long row with the inter-row spacing of 0.79 m and an alley space of 0.79 m. No irrigation was applied during the entire crop season as rainfalls were uniformly distributed for satisfactory plant growth. Conventional fertilizer, pest and weed control practices for growing field maize at Purdue University were followed.

### Phenotyping and data collection

The plant height and stalk width were measured in both wildtype (WT) and mutant (MT) siblings segregating in each family in the F_1_ populations of *Oy1*-*N1989*/*+*:B73 with the IBM, Syn10, and NIL populations. As described previously (Khangura *et al*. 2019a, 2019b), genetic backgrounds containing the *vey1^B73^* allele progressively suppresses the mutant phenotype and were tagged at V5-V7 stage when they were distinguishable from the wildtype siblings so that traits could be recorded at maturity when the genotypes are harder to distinguish. Trait measurements were performed on two to four WT and MT siblings that were picked at random. Plant height and stalk width were measured in cm and caliper units (Chaikam et al., 2011). Briefly, around three weeks after reproductive maturity, plant height was measured as the vertical distance from the soil line to the point of attachment of the flag leaf (abbreviated as FlHT), and ear height was measured as the vertical distance from the soil line to the top/primary ear (abbreviated as EaHT). Both FlHT and EaHT were recorded in cm. Additional height traits were calculated indirectly by subtracting EaHT from FlHT to get the distance between the top ear and flag leaf (abbreviated as Ea2Fl). Ratios (MT/WT) and differences (WT-MT) of these traits were also derived for each family. The EaHT were not measured in the IBM F_1_ populations, and consequently, any height trait derived from EaHT is missing in this population. Stalk widths (abbreviated as SW) were measured approximately three weeks after reproductive maturity using a Vernier caliper (Vernier 1631) at the middle of the first internode above the primary ear. The stalk width measurements were recorded as caliper units (c.u.), where 1 c.u. = 1/32 of an inch or ∼0.08 cm. The measurements of all the primary traits in two bi-parental and NIL populations are provided as **Tables S1-S3**.

Additional morphological features were quantified as follows. Tassel measurements were performed on wildtype and mutant siblings in the F_1_ progenies of 22 B73-like NIL and 13 Mo17-like NIL crossed with *Oy1-N1989/oy1*:B73 pollen-parents. Tassels were removed from mature plants at the point of flag leaf attachment and saved for measurements. The tassel traits measured included: Primary tassel branch number (TBN); Number of primary branches with at least one secondary branch (BN2); Secondary branch number (SecBN); Length of the tassel from point of flag leaf attachment to the tip (L1); Length of the main rachis of tassel (L2); Length of the branching zone (L3); Length of the topmost primary tassel branch (TopBLen); Length of the middle primary tassel branch (MidBLen); Length of the bottom primary tassel branch (LowBLen); Average of top, middle, and bottom primary tassel branch (AvgBLen). Tillering was assessed by counting the number of visible lateral primary branches one month after reproductive maturity. Mesocotyl measurements were done on seedlings grown under continuous red light (15-18 µmol m^-2^ s^-1^), far-red light (6-7 µmol m^-2^ s^-1^), white light (150 µmol m^-2^ s^-1^), and dark at 25° C for seven days at previously described peak wavelengths (Puthiyaveetil 2008; Macadlo *et al*. 2019).

### Genotypes and gene expression data

The genotypic and gene expression datasets used in this study are described previously in detail (Khangura *et al*. 2019a, 2019b). Briefly, the public marker data for IBM consists of 2,156 unique markers. The gene expression of the *oy1* locus was obtained from MaizeGDB. This public dataset consists of normalized read counts, expressed as reads per kilobase of transcript per million mapped reads (RPKM), of the maize genes from the transcriptome of shoot apices of 14 days old IBM seedlings (Li *et al*. 2013, 2018). The Syn10 genotypes were obtained from Liu et al., 2015. This dataset consists of SNP genotypes at 6611 positions (B73 RefGen v2). The BM-NIL carrying introgressions of the *vey1* region from opposite parents were selected for QTL validation from a set of previously described NIL (Eichten *et al*. 2011).

### Statistical analyses

Preliminary data exploration and descriptive statistics were calculated using JMP 13.0 (SAS Institute Inc. 2016). Trait means for each line were used as input phenotypes for QTL analyses. The QTL analyses were done using the R/QTL package version 1.40-8 (Broman et al., 2003) using single interval mapping (SIM) for all traits. The statistical thresholds to declare significant loci for each trait were computed by permutation testing using 1000 iterations (Churchill and Doerge 1994).

## Results

### *Oy1-N1989* reduces stalk width and is epistatic to *vey1*

Heterozygous *Oy1-N1989* plants (*Oy1-N1989/+*) have their eponymous oil-yellow color due to reduced chlorophyll content but are fertile and produce both ears and tassels (Khangura *et al*. 2019a). In the B73 inbred background, *Oy1-N1989*/+ plants germinate as yellow-green seedlings but the phenotype is progressively suppressed as they develop and are difficult to distinguish by color from wildtype plants at maturity (Khangura *et al*. 2019a). We previously described cis-acting expression polymorphisms at *oy1* distinguishing the maize inbred lines Mo17 and B73 and proposed that this expression polymorphism is the genetic basis of an *Oy1-N1989* modifier QTL *very oil yellow 1 (vey1)*. When the enhancing *vey1* allele from Mo17 is introgressed into the B73 background, *Oy1-N1989/+* plants with a *vey1^Mo17^* allele exhibited lower levels of chlorophyll and delayed flowering as compared to suppressed mutants with the *vey1^B73^* allele (Khangura *et al*. 2019a, 2019b). The further reduction in chlorophyll content in enhanced *Oy1-N1989/+* plants with a *vey1^Mo17^* allele increased the time to reproductive maturity and decreased CO_2_ assimilation and sugar contents in young leaves (Khangura *et al*. 2019b).

To determine if the reduced net CO_2_ assimilation, sugar content, or chlorophyll reduction affected by variation in *oy1* had effects on growth and development we measured multiple traits in materials with a range of phenotypic severities. We started with F_1_ crosses of B73 and Mo17 with and without the *Oy1*-*N1989* allele. As reported previously, chlorophyll measurements at two time-points (CCMI, ∼3-4 weeks after sowing; CCMII, ∼6-7 weeks after sowing) were lower in mutants in Mo17 x B73 F_1_ background compared to mutants in B73 background (**Table S4;** Khangura *et al*. 2019a). The heterotic Mo17 x B73 F_1_ wildtype hybrid plants had greater stalk diameters than the B73 plants (**Table S4**). The presence of an *Oy1-N1989* allele in the Mo17 x B73 F_1_ background eliminated the hybrid advantage producing mutant hybrids with narrower stalks than both wildtype and mutant B73 inbred plants. We hypothesized that the *vey1* allele in Mo17 was responsible for enhancing the effect of *Oy1-N1989* on stalk width in the B73 x Mo17 F_1_ hybrids. To test this, we measured stalk width in F_1_ populations generated by crossing *Oy1-N1989/+* to hundreds of recombinant inbred lines from the IBM and Syn10 populations. As expected, mutant chlorophyll values showed high positive correlations with mutant stalk width in both IBM and Syn10 F_1_ populations (**Tables S5 and S6**). Not surprisingly, as previously reported for chlorophyll content, CO_2_ assimilation, and reproductive maturity, the *vey1* locus modified stalk width in the *Oy1-N1989/+* mutant F_1_ siblings but not their wildtype individuals. The *vey1* locus was the largest effect QTL for stalk width in mutant siblings in both mapping populations explaining ∼43% and ∼68% phenotypic variation in IBM and Syn10 F_1_ populations, respectively (**Figure 1; Tables S7 and S8**). Assessment of stalk width to control for congenic wildtype sibling background using ratio only detected *vey1* in IBM, whereas in Syn10, an additional QTL was detected on chromosome 9. The regression using linked markers for this additional QTL (chr09.159.5) and *vey1* (chr10.94.5) in Syn10 did not detect any interaction between these two QTL for Ratio_SW (data not shown). No genes with previously described roles were present within a 400kb window surrounding the peak maxima on chromosome 9 (data not shown). These results formally extend the epistatic relationship of the *Oy1-N1989* allele and *vey1* locus to stalk width. The F_1_ mutant plants that inherited the suppressor *vey1^B73^* allele were thicker than the mutant plants harboring enhancing *vey1^Mo17^* allele. This demonstrated that stalk width responded similarly to chlorophyll content, reproductive maturity, and CO_2_ assimilation. Just as was the case for chlorophyll content, *vey1* was the only QTL detected for mutant stalk width in both populations.

**Figure 1.**
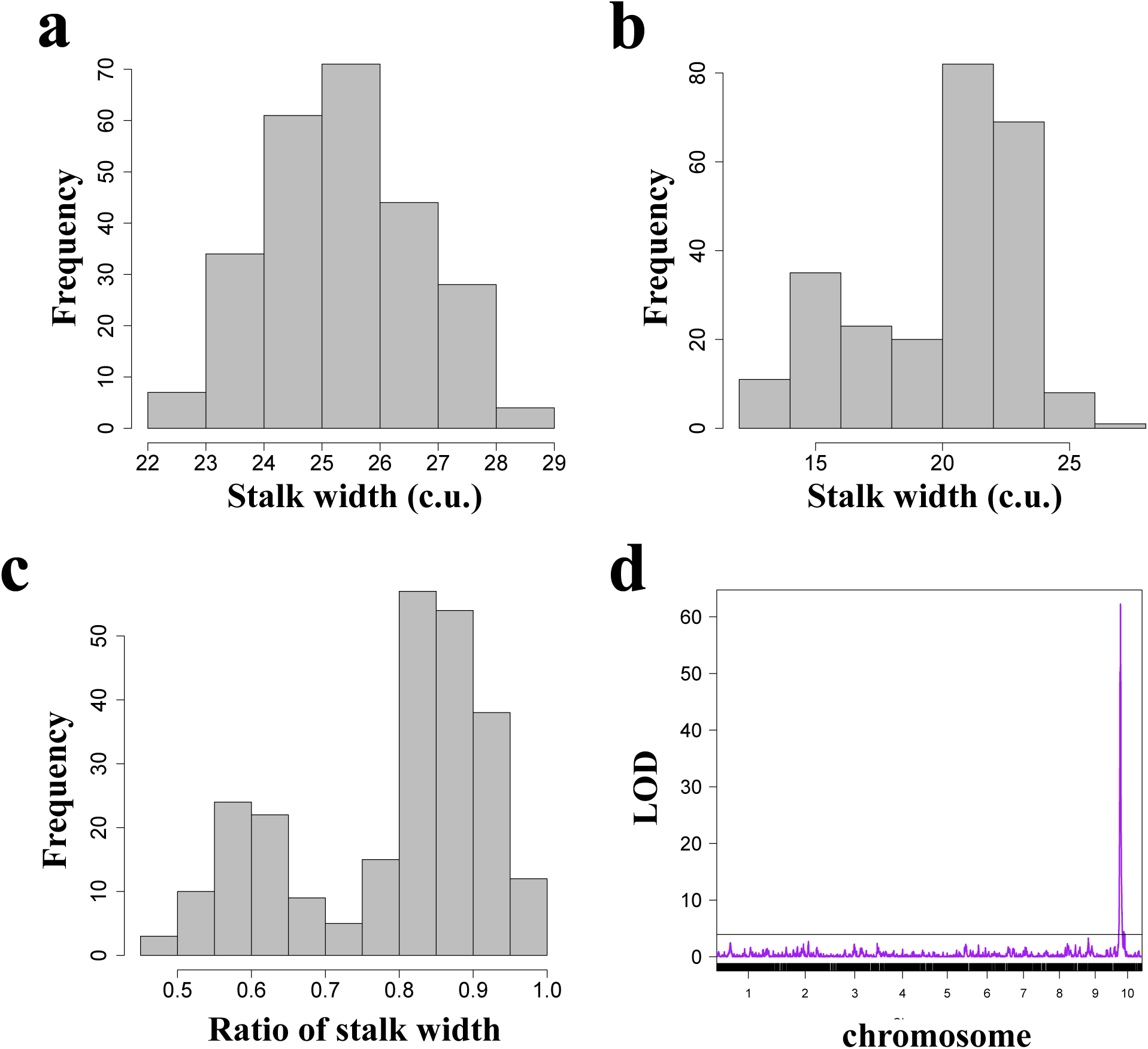
The trait distributions and QTL plot of stalk width in the Syn10 x *Oy1-N1989*/+:B73 F_1_ population. The trait distribution of (a) wildtype siblings, (b) mutant siblings, and (c) ratio of mutant/wildtype. (d) The genome-wide QTL result of mutant stalk width. The stalk width measurements are expressed as caliper units (c.u.), with 1 c.u. = 1/32 of an inch or ∼0.08 cm.

The stalk widths of wildtype siblings in both populations were also variable and under genetic control. The localization of small effect QTL are often imprecise, but candidate genes within a window bound by a 2-LOD decrease in the QTL significance were tabulated for each detected QTL (**Tables S7 and S8**). QTL-candidate genes were assessed for roles in growth regulation and marked for eQTL using data from a previous study of the IBM population (Li *et al*. 2013, 2018). Three QTL, one in the IBM and two in the Syn10 F_1_ population, were detected on chromosome 1 for wildtype stalk widths (**Tables S7 and S8**). There was no overlap of the 2-LOD QTL intervals between IBM and Syn10 for stalk width of the wildtype siblings. Moreover, the Mo17 allele at the QTL detected in the IBM increased wildtype stalk width, whereas both Syn10 QTL showed increase in stalk width with B73 alleles, making an imprecise localization of the same QTL detected in both populations on chromosome 1 unlikely. The QTL in IBM is a moderate effect QTL with an estimated contribution of 13% of the phenotypic variation (**Tables S7**), but such estimates should be regarded as upper estimates (Beavis 1994). The 2-LOD window at this QTL contains a candidate gene in the ß-expansin family (*expb4;* B73 v4: Zm00001d033231), whose ortholog in rice have been implicated to increase stem elongation in response to gibberellic acid (Lee and Kende 2001). *EXPB4* transcript abundance was influenced by cis and trans eQTL with B73 alleles increasing expression in all cases, making a cis-acting eQTL a possibility for this candidate gene. No other QTL overlapped genes with previously described roles in stem architecture (data not shown).

Confirmation of the impact of the *vey1* locus on stalk width was provided by crosses between *Oy1-N1989*/+ mutants and near-isogenic lines (NIL) with introgressed *vey1* alleles. In all NIL F_1_ families, *Oy1-N1989/+* mutants were thinner than their wildtype siblings (**Table S3**). In F_1_ crosses of NIL from the B73 and Mo17 isogenic backgrounds to *Oy1-N1989*/+, the allele at *vey1* had no discernable impact on the stem width of the wildtype siblings (**Figures 2a** and **S2**). In both B73 and Mo17 x B73 F_1_ isogenic backgrounds inheritance of the *vey1^Mo17^* allele enhanced the *Oy1-N1989/+* phenotype leading to thinner plants than isogenic siblings with the suppressing *vey1^B73^* allele (**Figures 2a** and **S2**). Thus, *Oy1-N1989* controlled stalk width and was epistatic to the *vey1* QTL, consistent with the previously reported effects and interactions of these alleles on chlorophyll content, reproductive maturity, and CO_2_ assimilation (Khangura *et al*. 2019a, 2019b).

**Figure 2.**
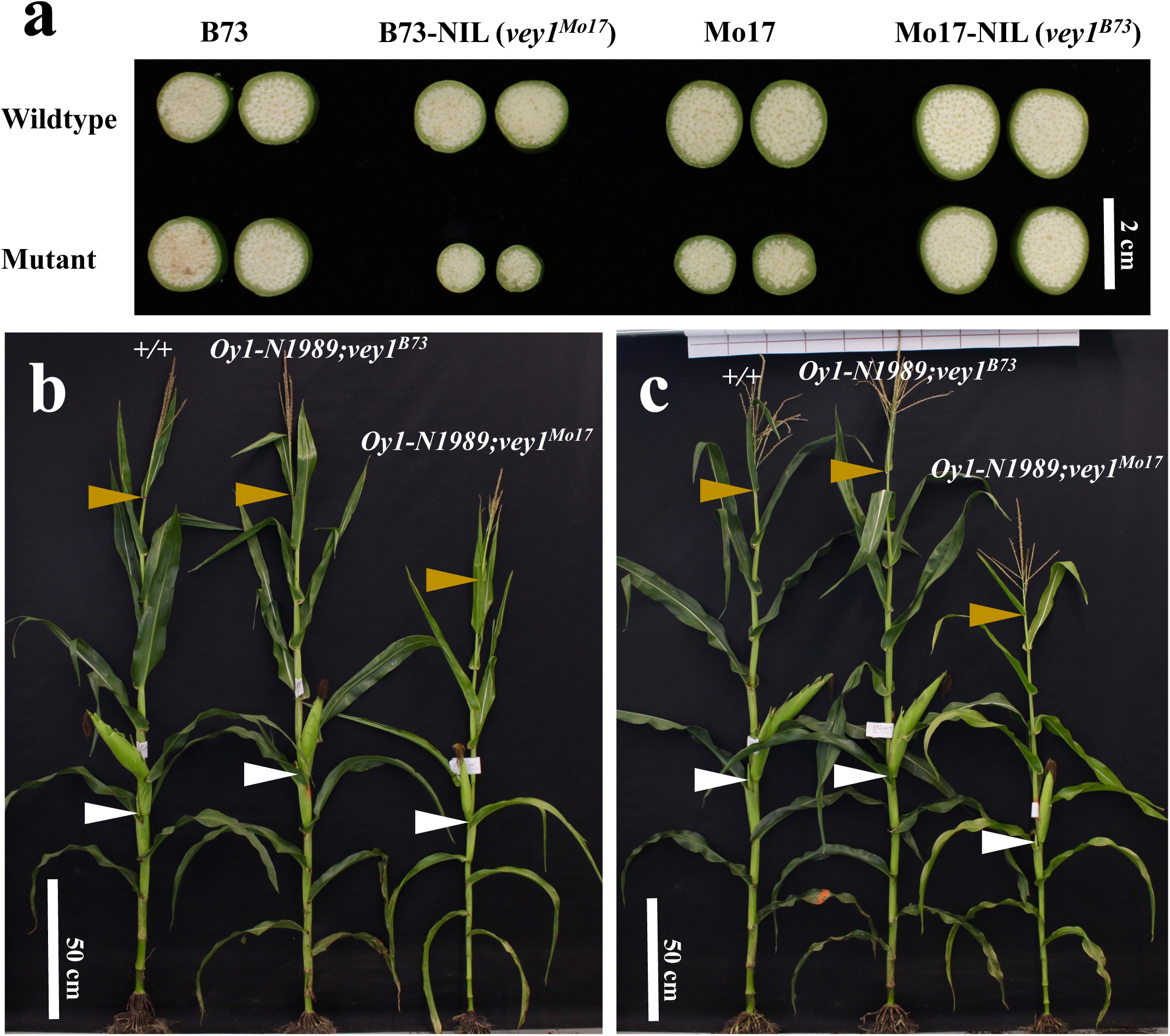
The effect of *vey1* alleles on stalk width and height in mutant and wildtype individuals in isogenic backgrounds. (a) The representative cross-sections from the middle of one internode above the top ear at maturity in the same material. (b-c) The representative wildtype and mutant plants at maturity with both *vey1^B73^* and *vey1^Mo17^* alleles in isogenic (b) B73 NIL x *Oy1-N1989/+*:B73, and (c) Mo17 NIL x *Oy1-N1989/+*:B73 crosses. The golden arrows indicate a flag leaf node, and the white arrows indicate the top or primary ear node

### Plant height in *Oy1-N1989* shows a non-linear effect direction dependent on the *vey1* allele

In addition to the previously described delay in reproductive maturity of *Oy1-N1989*/+ mutants in the B73 background (Khangura *et al*. 2019b), mutant plants reached or surpassed the total plant height (FlHT) and ear height (EaHT) of their congenic B73 wildtype siblings (**Figure 2b** **and Table S4**). In the parental crosses, wildtype Mo17 x B73 F_1_ hybrid plants had greater plant height and ear height than the B73 inbreds (**Table S4**). Not only were *Oy1-N1989*/+ mutants in the Mo17 x B73 F_1_ hybrid not taller than wildtype hybrids, they showed a significant reduction in ear height, but not plant height, compared to the wildtype siblings.

We measured height traits in the RIL x *Oy1-N1989*/+ F_1_ populations to determine what effect the *vey1* modifier would have on plant height in the *Oy1-N1989*/+ mutants. First, we measured total plant height (FlHT) in the IBM and Syn10 F_1_ populations. Just as we previously reported for chlorophyll content (Khangura *et al*. 2019a), flowering time, CO_2_ assimilation (Khangura *et al*. 2019b), and stalk width (**Figure 1**), linkage mapping detected a QTL at *vey1* for plant height in *Oy1-N1989*/+ mutant individuals but not wildtype siblings (**Figure 3; Tables S7 and S8**). On average, F_1_ mutant plants that inherited the *Oy1-N1989-*supressing *vey1^B73^* allele, were taller than *Oy1-N1989/+* mutant plants harboring the *vey1^Mo17^* allele (**Figure 4; Tables S7 and S8**). This effect was clear in both IBM and Syn10 F_1_ populations. In addition, the average height of mutant plants that inherited the *vey1^B73^* allele was significantly greater than their much greener wildtype siblings (**Figure 4**). In both populations, mutant plants that inherited the *vey1^Mo17^* allele were shorter than their wildtype siblings. It would appear that the relationship between the disruption of chlorophyll biosynthesis and final plant height are complicated and non-linear. As a result, *Oy1-N1989/+* mutants can either be taller or shorter than their wildtype siblings with the direction of the deviation conditioned by the *vey1* allele.

**Figure 3.**
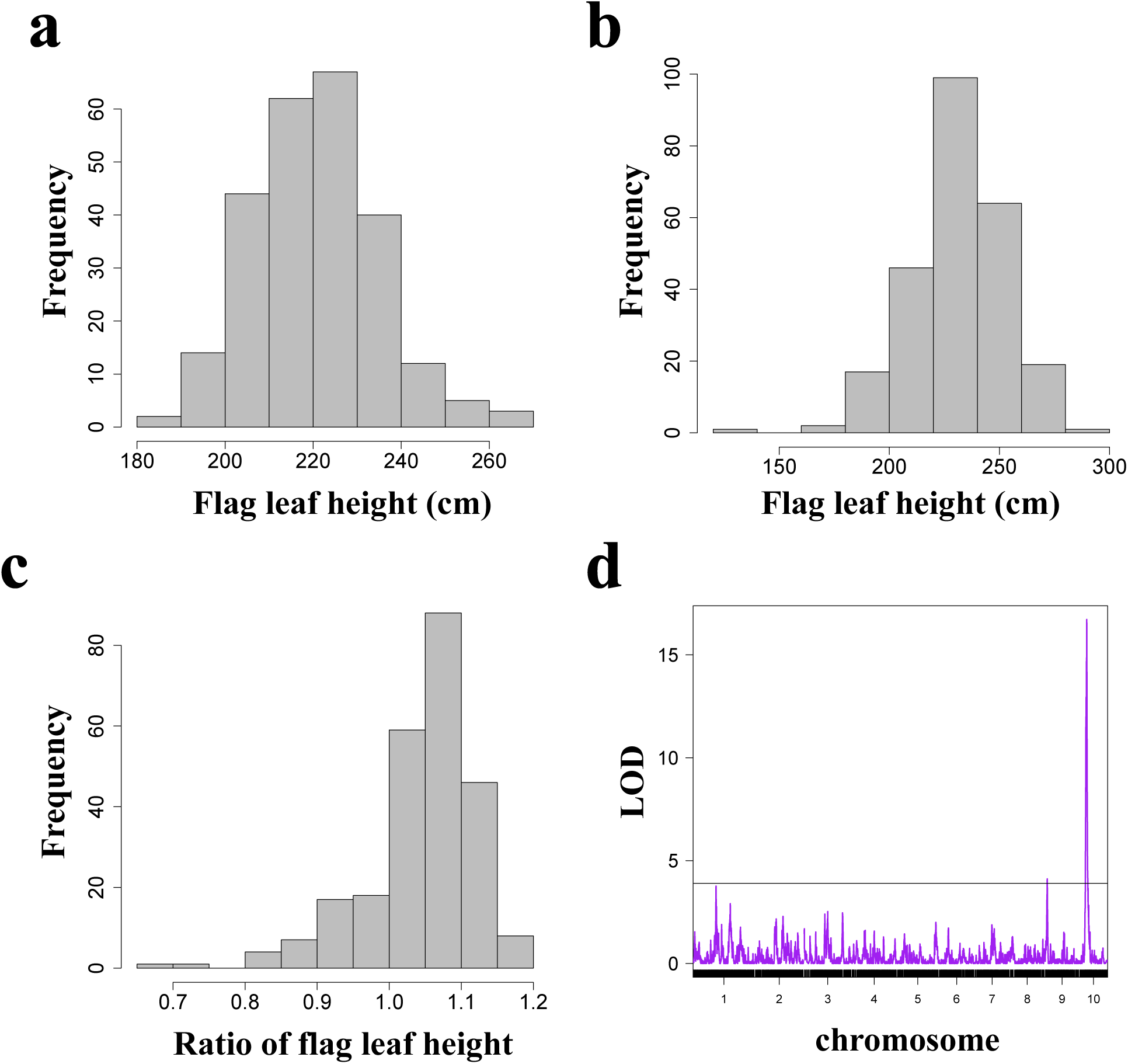
The trait distributions and QTL plot of plant height variation in the Syn10 x *Oy1-N1989*/+:B73 F_1_ population. The trait distribution of (a) wildtype siblings, (b) mutant siblings, and (c) mutant/wildtype. (d) The genome-wide QTL result of mutant plant height.

**Figure 4.**
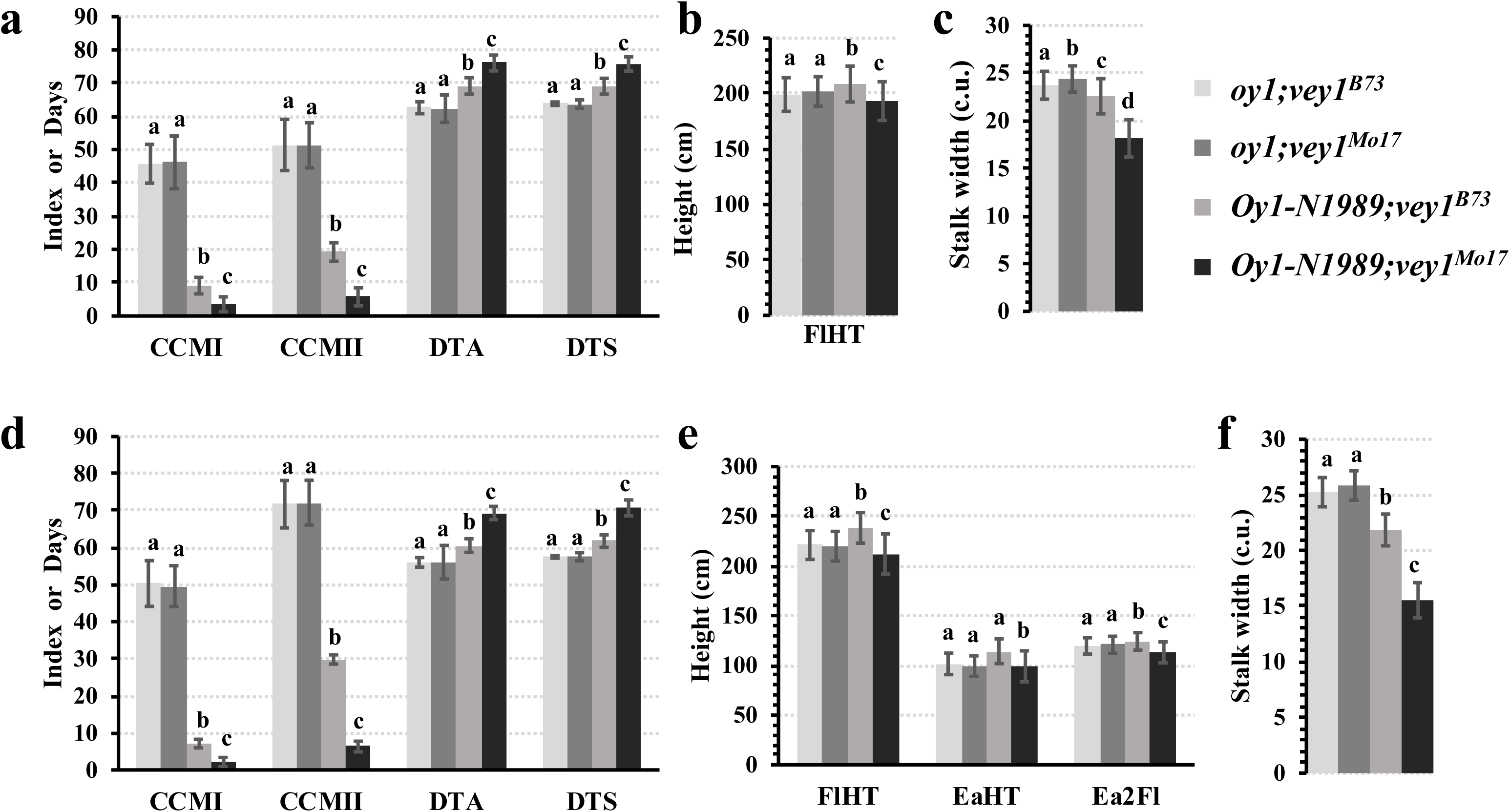
The effect of *vey1* alleles on various physiological and developmental traits in the (a-c) IBM x *Oy1-N1989/+*:B73 and (d-f) Syn10 x *Oy1-N1989/+*:B73 F_1_ hybrid populations. The chlorophyll content measurements (CCMI: early season; CCMII: late season) and time to reproductive maturity (DTA: days to anthesis; and DTS: days to silking) are depicted in panel a and d (data reanalyzed from Khangura *et al*., 2019a and 2019b); height in panel b and e; stalk width in panel c and f. The statistical significance was determined using ANOVA, and the connecting letter report indicates mean comparisons using Tukey’s HSD at p<0.05.

In the Syn10 F_1_ crosses, we also measured the height of the primary ear node (EaHT) which allowed us to obtain distance between primary ear node and flag leaf (Ea2Fl). Similar to the results for total plant height, *vey1^Mo17^* resulted in reduced EaHT and Ea2Fl in Syn10 F_1_ plants and *vey1^B73^* resulted in plants with greater EaHT and Ea2Fl than wildtype controls. Thus, plant heights of chlorotic *Oy1-N1989*/+ mutants can be either increased or decreased by alternate alleles of *vey1*. This allele-specific interaction between *vey1* and *Oy1-N1989*, with opposing effects on mutant plant height for *vey1^B73^* and *vey1^Mo17^* as compared to the wildtypes, is inconsistent with the simple interpretation that plant height in the *Oy1-N9189* mutants is conditioned solely by reduced growth resulting from less carbon assimilation in the *Oy1-N9189* mutants.

A number of smaller effect QTL were also identified for plant height in both IBM and Syn10 F_1_ populations (**Tables S7 and S8**). Using the same approach described above for stalk width, candidate genes were sought with a 2-LOD window of QTL that were not linked to *vey1* and affected height. A QTL for wildtype FlHT was detected on chromosome 3 in the Syn10 F_1_ population (**Table S8**) at a similar position to a previously reported plant height and flowering time QTL in both Syn10 and IBM inbred lines (Liu *et al*. 2015). This QTL overlaps the recently described flowering time regulator *zmmads69* (B73 v4: Zm00001d042315; Liang *et al*. 2019) encoding a QTL detected between maize and teosinte (Liang *et al*. 2019), and in two maize association panels (Hirsch *et al*. 2014; Lin *et al*. 2017). Over-expression of *zmmads69* hastens flowering time in maize and reduces plant height in all cases (Liang *et al*. 2019). Consistent with *zmmads69* encoding this QTL and the direction of the effect of the QTL on height (**Table S8**), a strong cis-eQTL encodes increased accumulation of mRNA from the Mo17 allele of *zmmads69* (LOD = 44.17, additive effect = -3.84, R^2^ = 0.76; Table S3 from Li *et al*. 2013; please note that the effect direction was reversed due to the correction in Li *et al*. 2018). We tested the mutant siblings for an effect of *zmmads69* on flowering time and for interaction with *vey1* using the data of Khangura *et al*. 2019b. The marker associated with reduced wildtype FlHT linked to *zmmads69* (chr03.1591) was significant in a single marker regression test for an effect on flowering time in the wildtype (LOD = 3.04, additive effect = 0.57, R^2^ = 0.05, P-value = 0.0002) and mutant siblings (LOD = 0.86, additive effect = 1.13, R^2^ = 0.02, P-value = 0.047) but had no effect on the ratio or difference in flowering time between mutant and wildtype siblings (data not shown). There was no interaction between *zmmads69* and *vey1* for days to anthesis in mutant siblings (data not shown) indicating that this QTL is not chlorophyll-dependent. Interaction between *zmmads69* and *vey1* for mutant height traits (FlHT, EaHT, and Ea2Fl) were only significant for mutant Ea2Fl (additive effect = -6.58, R^2^ = 0.02, P-value = 0.03) but not in the normalized Ea2Fl (both ratio and difference). To test if two flowering time QTLs, *vey1* and *other oy1 flowering time locus1* (*oof1*), from mutant (Khangura *et al*. 2019b) also result in plant height differences, we performed a regression test using the main and interaction effects of these QTLs on mutant FlHT. There was no effect of *oof1* or *oof1* x *vey1* interaction on mutant FlHT (data not shown).

Multiple FlHT QTL were identified on chromosome 9 in both Syn10 and IBM F_1_ populations (Tables S7 and S8). In Syn10 mutant F_1_ plants we detected a FlHT QTL, centered at 3.2 Mbp, at the top of chromosome 9 close to the position of height QTL for wildtype plants from B73 x Mo17 crosses detected in multiple previous studies (Beavis *et al*. 1991, 1994; Austin *et al*. 2001; Liu *et al*. 2015). Others have noted that this QTL colocalizes with the *dwarf plant3* (*d3*) gene (Beavis *et al*. 1991; Wang *et al*. 2006, 2018) encoding Ent-kaurenoic acid oxidase (B73 v4 Zm00001d045563; Winker and Helentjaris 1995). There is a strong cis-eQTL in the IBM transcriptomic data that accounts for 63% of the variation in DWARF PLANT3 mRNA accumulation (Li *et al*. 2013). Recently, it was revealed that this eQTL study of the IBM population (Li *et al*. 2013) had reversed the signs of the effects on gene expression (Li *et al*. 2018). As a result of this correction, it is clear that *dwarf plant3* cannot encode this QTL as the gene expression is lower from the Mo17 allele, and the linked marker with Mo17 allele condition an increase in plant height (**Table S8**). The narrow 2-LOD window around the peak of 3.2 Mbp for our localization of this QTL also overlapped with the *pin1a* gene (B73 v4: Zm00001d044812; Forestan *et al*. 2012), encoding an auxin efflux carrier. No eQTL was observed for *pin1a* suggesting a coding sequence polymorphism would be required for this to be a plausible candidate gene. A second QTL on chromosome 9, with a peak at 12 Mbp and unlinked to any other QTL, was detected for wildtype Ea2Fl in the Syn10 F_1_ population. A third QTL on chromosome 9 controlled multiple wildtype and mutant height traits with peak at 106 and 118 Mbp, respectively. The mutant FlHT QTL centered at 118 Mbp overlapped with *zmgras45* (B73 v4: Zm00001d047123; Guo *et al*. 2017), orthologs of which have been previously shown to influence gibberellin-mediated regulation of chlorophyll biosynthesis (Ma *et al*. 2014). No eQTL was detected for *zmgras45* in the previous study of IBM shoot apices (Li *et al*. 2013, 2018).

Of the remaining QTL, only one had a 2-LOD window that overlapped a gene known to impact plant height. In the Syn10 F_1_ population, a QTL on chromosome 1 with a peak at 82 Mbp was detected for FlHT in both wildtype and mutant siblings (**Table S8**). The 2-LOD window for this locus overlaps a MATE transporter (MATE; B73 v4: Zm00001d029677), which is orthologous to *A. thaliana* genes known to encode dominant alleles affecting hypocotyl elongation (Wang *et al*. 2015) and facilitating ABA transport (Zhang *et al*. 2014). No significant eQTL were described for this MATE in the previously published IBM eQTL study (Li *et al*. 2013, 2018).

### Single locus recapitulation and validation of the non-linear relationship between plant height and chlorophyll level

To test the single-locus effect of *vey1* on plant height of the *Oy1-N1989* mutant, we crossed *Oy1-N1989/+* mutants to near-isogenic lines (NIL) in B73 background with *vey1^Mo17^* allele introgressions. The F_1_ progenies from these crosses segregated for *Oy1-N1989* mutants with alternative *vey1* alleles (*vey1^B73^* and *vey1^Mo17^*). Supporting a simple explanation of one-locus determination, in the isogenic B73 background the total plant height of *Oy1-N1989/+* mutants with a *vey1^B73^* allele was greater than both wildtype siblings and *Oy1-N1989*/+ mutants with a *vey1^Mo17^* allele (**Figures 2** and **5**). Similarly, when *vey1^Mo17^* enhanced the *Oy1-N1989*/+ phenotype, plants were shorter than all other genotypes. Thus, we were able to recapitulate the effect of *vey1* on mutant plant height observed in the RIL and DH population using a single-locus introgression in an isogenic inbred background.

**Figure 5.**
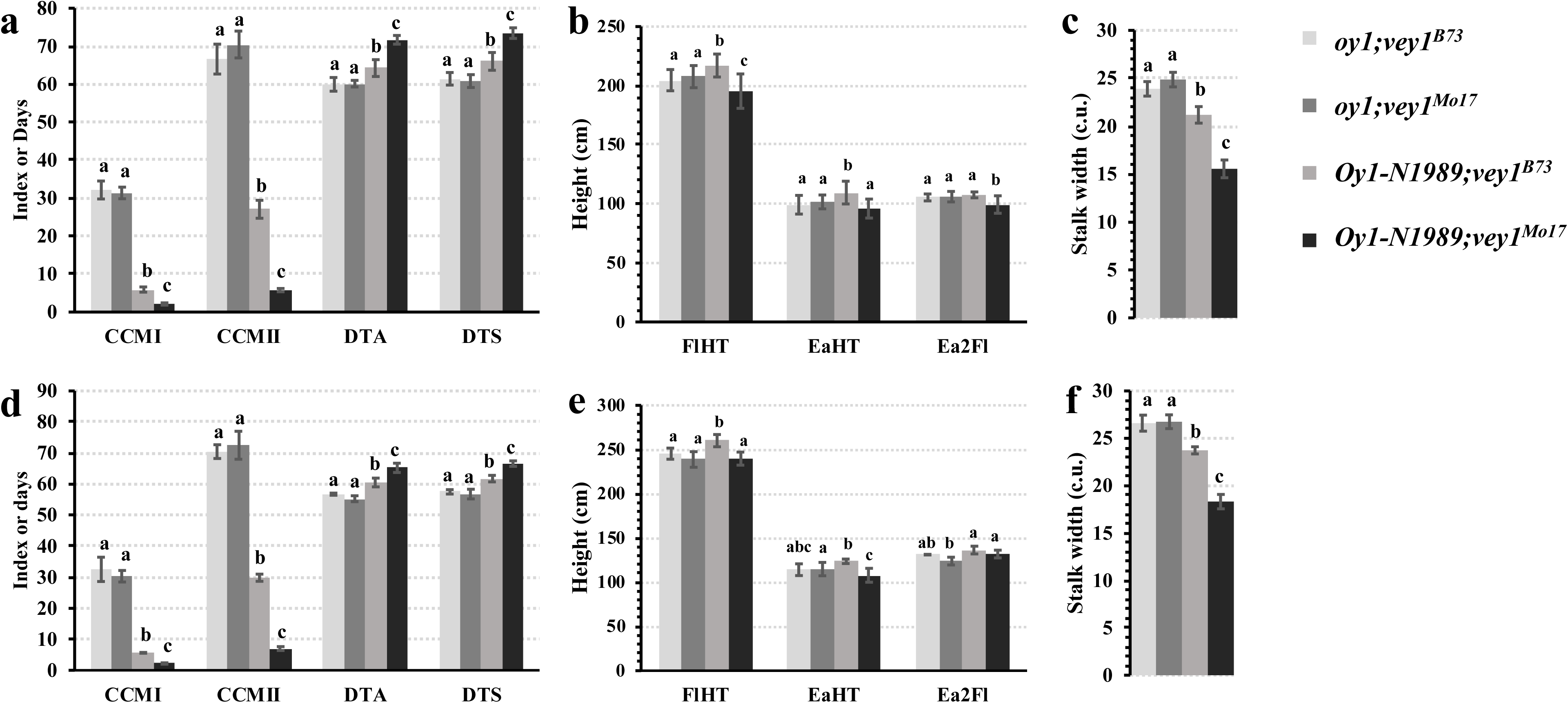
The effect of *vey1* alleles on various physiological and developmental traits in the (a-c) B73-like NILs x *Oy1-N1989/+*:B73 and (d-f) Mo17-like NILs x *Oy1-N1989/+*:B73 F_1_ hybrid populations. The chlorophyll content measurements (CCMI: early season; CCMII: late season) and time to reproductive maturity (DTA: days to anthesis; and DTS: days to silking) are depicted in panel a and d (data reanalyzed from Khangura *et al*. 2019a, 2019b); height in panel b and e; stalk width in panel c and f. The statistical significance was determined using ANOVA, and the connecting letter report indicates mean comparisons using Tukey’s HSD at p<0.05.

The B73 NIL removed hybridity from our tests, and we considered that some of the variation in the *Oy1-N1989* x *vey1* interaction effects on plant height could be driven by overall plant vigor and hybridity. For instance, in both IBM and Syn10 crosses, the *vey1^Mo17^* allele decreased plant height (**Figure 4B** **and 4E**), but we did not see such reduction in height in the original B73 x Mo17 F_1_ materials that also contained *vey1^Mo17^* allele (**Table S4**). Each RIL and DH is ∼50% Mo17 while the *Oy1-N1989* mutant is in a B73 inbred background resulting in F_1_ progeny that were hybrid at a random collection of 50% of the loci contributing to heterosis in this cross. To perform a test of the effect of *vey1* in an isogenic hybrid background, we crossed *Oy1-N1989/+*:B73 to NIL derived using Mo17 as the recurrent parent with introgressions of alternative alleles at *vey1*. This test maintains hybridity at most unlinked loci between B73 and Mo17, unlike the RIL crosses which retained hybridity for only approximately half of the genome. The mutant F_1_ progenies derived from Mo17 NIL containing the *vey1^B73^* allele were significantly taller than the wildtype siblings (**Figures 2** and **5**), just as observed in the IBM, Syn10, and isogenic B73 NIL F_1_ progenies (**Table S4;** **Figures 2**, **4**, and **5**). However, the mutant F_1_ progeny derived from Mo17 NIL that inherited the *vey1^Mo17^* allele showed no discernable difference to total plant height as compared to wildtype siblings, but ear height decreased and a compensatory increase in the height between ear and flag leaf was observed (**Figure 5**). Thus, *vey1^B73^* consistently increased plant height of the *Oy1-N1989/+* plants over wildtype siblings in isogenic B73 background, across a population of recombinant inbred lines crossed to B73, and in B73 x Mo17 F_1_ NIL hybrids, whereas, the *vey1^Mo17^* allele decreased mutant plant heights when compared to wildtype siblings in all materials except the hybrid Mo17 x B73 F_1_ background.

### Expression level variation at *oy1* in IBM correlates with stalk width and height in mutants

We previously proposed *vey1* encodes a cis-acting expression polymorphism in the regulatory sequence at *oy1* (Khangura *et al*. 2019a, 2019b). Our previous reanalysis of transcript data derived from shoot apices from IBM (Li *et al*. 2013, 2018), a maize association panel (Kremling *et al*. 2018), and our allele-specific expression assays in NIL demonstrated that the *vey1^B73^* allele was linked to higher expression of wildtype OY1 transcript (Khangura *et al*. 2019b). Based on this linkage, we expect stalk width and height variation in mutant plants to also be correlated with OY1 mRNA accumulation. Stalk width and plant height in mutant IBM F_1_ progenies showed significant linear correlation with OY1 transcript abundance in the shoot apices of the IBM inbred lines (**Figure 6**). The OY1 transcript variation in IBM did not correlate with stalk width and plant height in the wildtype siblings. These results are consistent with *vey1* encoding a cis-regulatory polymorphism controlling OY1 as the genetic basis of stalk width and height traits in *Oy1-N1989*/*+* mutants.

**Figure 6.**
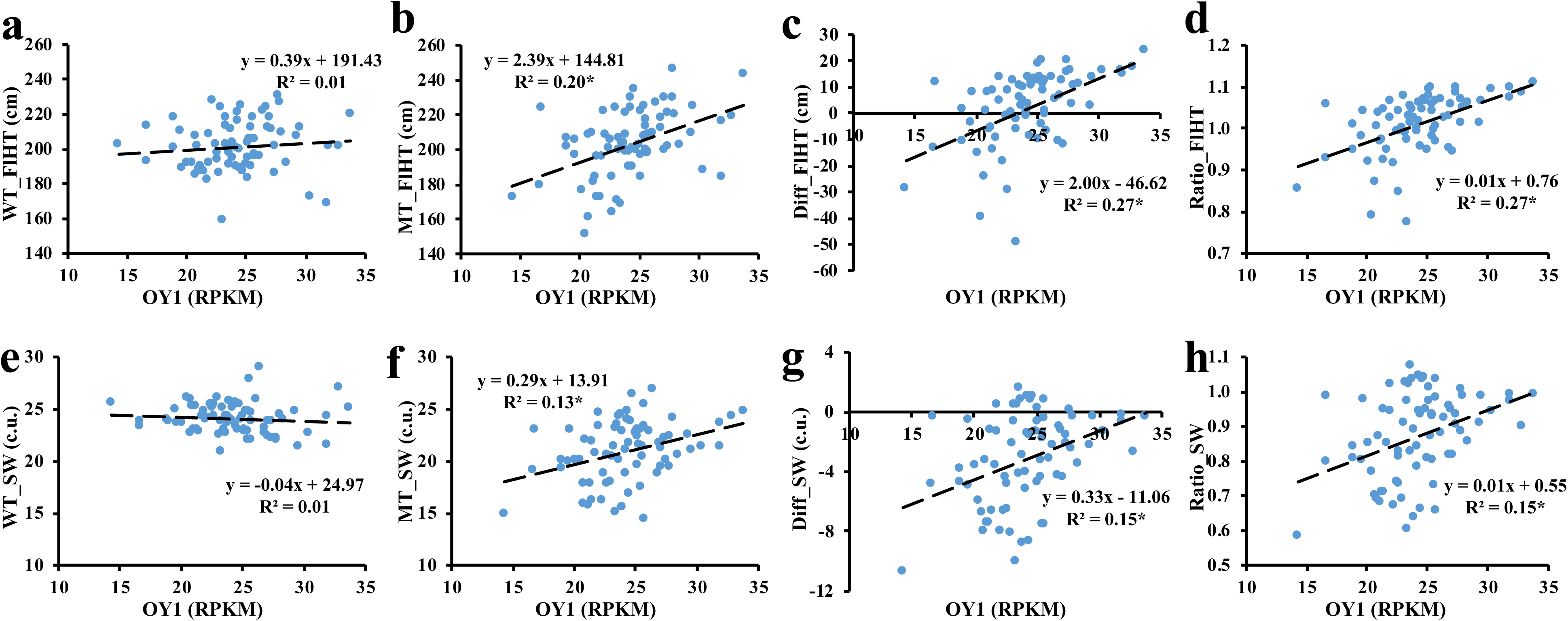
The linear regression of normalized (expressed as RPKM) read counts of OY1 transcripts (X-axis) in IBM seedlings (n = 74) with (a-d) plant height, and (e-h) stalk width (SW) measurements of wildtype (WT) and mutant (MT) siblings in IBM x *Oy1-N1989/+*:B73 F_1_ population. The equation for the linear fit and R^2^ value of the model is listed on each plot. The asterisk at the end of the R^2^ value indicates statistical significance at p<0.0001. Panel c, d, g, and h contain derived plant height and stalk width traits (Diff: MT-WT; Ratio: MT/WT). Check methods for detail.

If the effect of *vey1* on *Oy1-N1989*/*+* mutants is the result of rescuing chlorophyll biosynthesis, rather than some novel function of the protein encoded by *Oy1-N1989*, then chlorophyll abundance should correlate better with gross morphology than OY1 transcript abundance. Both CCM measures (CCMI and CCMII) vary within each *vey1* genotypic class. CCM increases even within the RIL containing *vey1^Mo17^* and one containing *vey1^B73^* allele (**Figures S3-S7**). If the effects on development are mediated by a loss of net CO_2_ capture and plant energy status due to reduction in chlorophyll, then the variation in CCM levels in *Oy1-N1989* mutants due to phenomena other than *vey1* should result in corresponding fluctuation in phenotype.

To explore the relationship between chlorophyll abundance variation affected by *Oy1-N1989* and modifiers and development, we explored the phenotypic correlations between the previously measured and described CCM data (Khangura *et al*. 2019a), the reproductive maturity date (Khangura *et al*. 2019b), plant height and stalk width measurements detailed in this study. As the effect of *vey1* is only visible in the presence of the *Oy1-N1989* allele for all traits, we only used difference of congenic wildtype and mutant siblings (MT-WT) to study these variations with respect to chlorophyll. Mutant stalk widths reduced (increase in difference) with a decrease in CCM within both *vey1^Mo17^* and *vey1^B73^* genotypes in IBM and Syn10 F_1_ populations (**Figure S3**). Consistent with the effects on net CO_2_ capture (Khangura *et al*. 2019b), the slope of the regression line and therefore the fit was greater for the mutants with *vey1^Mo17^* allele. Thus, the variation in chlorophyll content among individuals within each *vey1* genotype was sufficient to affect stalk width.

Similarly, the reduction in total plant height was also suppressed in mutants with *vey1^Mo17^* allele that accumulated relatively higher chlorophyll in both IBM (R^2^ = 0.18, P-value = <0.0001) and Syn10 (R^2^ = 0.09, P-value = 0.008) F_1_ populations (**Figure S4**). There was no discernable impact of a change in chlorophyll content on plant height in the *vey1^B73^* mutant F_1_ siblings (**Figure S4**). The effect of chlorophyll variation on ear height was only significant between the later mutant chlorophyll estimate (CCMII) in IBM with *vey1^Mo17^* allele and ear height, although a similar trend was observed for Syn10 F_1_ progenies inheriting *vey1^Mo17^* allele (**Figure S5**). As expected for a simple relationship, as mutant CCM values increased the mutant plants matured earlier as determined by days to anthesis (DTA) in both *vey1* classes in both IBM and Syn10 (**Figures S6 and S7**). Regardless of the reason for these effects, the increase in plant height observed in *Oy1-N1989* mutants with *vey1^B73^* allele remains the outlier for the relationship and effect of *vey1* alleles on the *Oy1-N1989* mutant phenotype.

### Seedling etiolation in far-red light is not altered in *Oy1-N1989* mutants

The non-linearity of the effect of *Oy1-N1989* and *vey1* on plant height raised the possibility that disrupting CHLI alters light signaling in maize. The shared biosynthetic origins of chlorophyll and bilin chromophores required for phytochrome assembly (Frankenberg and Lagarias 2012) was explored as a possible source of increase in total plant height observed in *Oy1-N1989/+* mutants modified by *vey1^B73^*. Low R/FR ratio is indicative of shade in the plant canopy and elicits a shade avoidance syndrome via phytochrome signaling (Chen *et al*. 2004; Pierik and De Wit 2014; Roig-Villanova and Martínez-García 2016; Ortiz-Alcaide *et al*. 2019). If a mutation in *oy1* perturbs tetrapyrrole biosynthesis, a reduction in the pool of functional phytochrome should increase mesocotyl length and plant height and reduce the number of days to reproductive maturity as had been reported previously in the *elongated mesocotyl1* (*elm1*) maize mutant which is defective in chromophore synthesis and has reduced phytochrome levels (Sawers *et al*. 2002). To test whether the range MgChl defects affected by the *Oy1-N1989* and *vey1* interactions influence the phytochrome pool, we grew plants in a variety of light conditions that stimulate phytochrome mediated elongation.

We grew near-isogenic *Oy1-N1989* mutant seedlings with segregating *vey1* alleles under four light conditions (continuous dark, white, red and far-red light) and measured mesocotyl length. The *elm1* mutant was used as a positive control for these growth conditions (Sawers *et al*. 2002, 2004). As expected, the *elm1* mutants showed significantly longer mesocotyls compared to wildtype in all light conditions, and the stimulatory effect of red and shade-mimicking far-red light on mesocotyl length in *elm1* was evident (**Figure 7**). Under all light conditions tested, no statistically significant differences were observed in mesocotyl length between wildtype and *Oy1-N1989* mutants with either *vey1^Mo17^* or *vey1^B73^* allele. These results indicate no, or very limited, impact *Oy1-N1989* on light-mediated growth responses consistent with independent regulation of the tetrapyrrole pathway branches leading to chlorophyll and chromophore biosynthesis.

**Figure 7.**
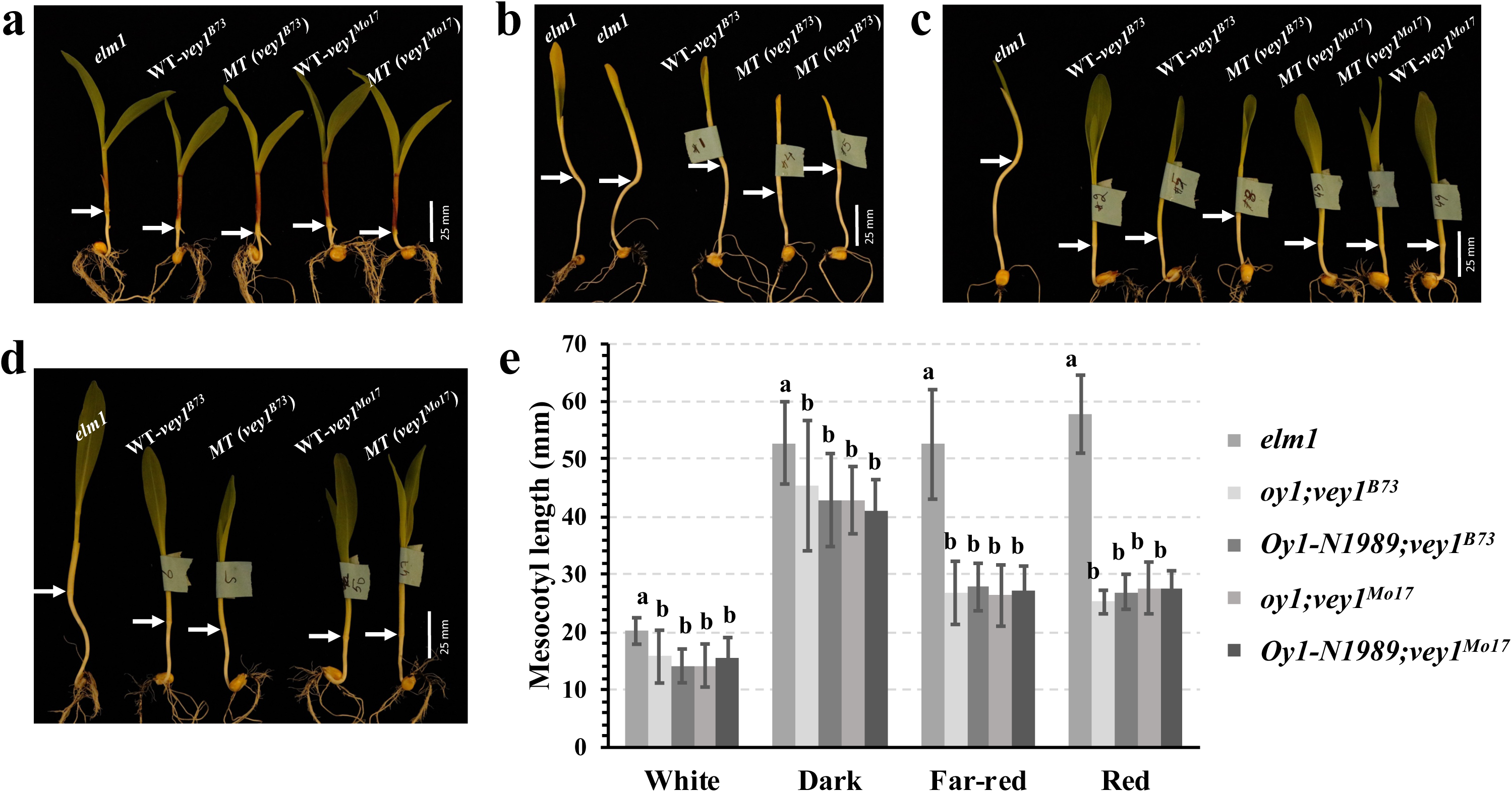
The seedling phenotypes of *elm1,* wildtype (WT) and *Oy1-N1989*/+ mutants (MT) with both *vey1* alleles (*vey1^B73^* and *vey1^Mo17^*) after 7 days of growth under continuous (a) white light, (b) no light/dark, (c) red light, and (d) far-red light conditions. The white arrows in panel a-d indicate boundary between mesocotyl and the first internode. (e) The quantification (mean ± S.D.) of various genotypic groups in all four growth conditions at 7 days. The connecting letter report indicates statistical significance determined using the student’s t-test at p<0.05. The sample size is 8-19 seedlings per genotype in each growth condition.

### Tassel branching is reduced in severe *Oy1-N1989* mutants

A recessive mutant allele at *oy1* was previously shown to reduce the tassel branch number in maize (Brewbaker 2015). The availability of near-isogenic materials with segregating *vey1* alleles prompted us to test whether the semi-dominant *Oy1-N1989* allele and its interaction with *vey1* alters tassel branching and development. Using the F_1_ populations derived from crosses of both B73-like and Mo17-like NIL with *Oy1-N1989/+*:B73, we performed a morphometric analysis of the tassels collected from mutant and wildtype siblings of each F_1_ family. Tassel growth (measured as the length of the total tassel, branching zone, and central spike) and development (primary and secondary branch number) was not affected in *Oy1-N1989/+* mutants modified by the suppressing *vey1^B73^* allele in both B73 and Mo17 NIL backgrounds (**Figure 8** **and Table S9**). The *Oy1-N1989/+* mutants carrying the enhancing *vey1^Mo17^* allele had reduced tassel branch numbers and tassel length. Mutants derived from crosses of *Oy1-N1989/+*:B73 with B73 and Mo17 NIL carrying *vey1^Mo1^*^7^ introgression displayed a reduction in the primary tassel branch number by ∼47% and ∼23%, respectively. Similarly, the overall length of the tassels was reduced by ∼39% and ∼18% in *Oy1-N1989* mutants in the F_1_ generation with B73 and Mo17 NIL with *vey1^Mo1^*^7^ introgression, respectively. These changes in tassel architecture are consistent with the report of Brewbaker (Brewbaker 2015) where ∼30% reduction in tassel branching was observed.

**Figure 8.**
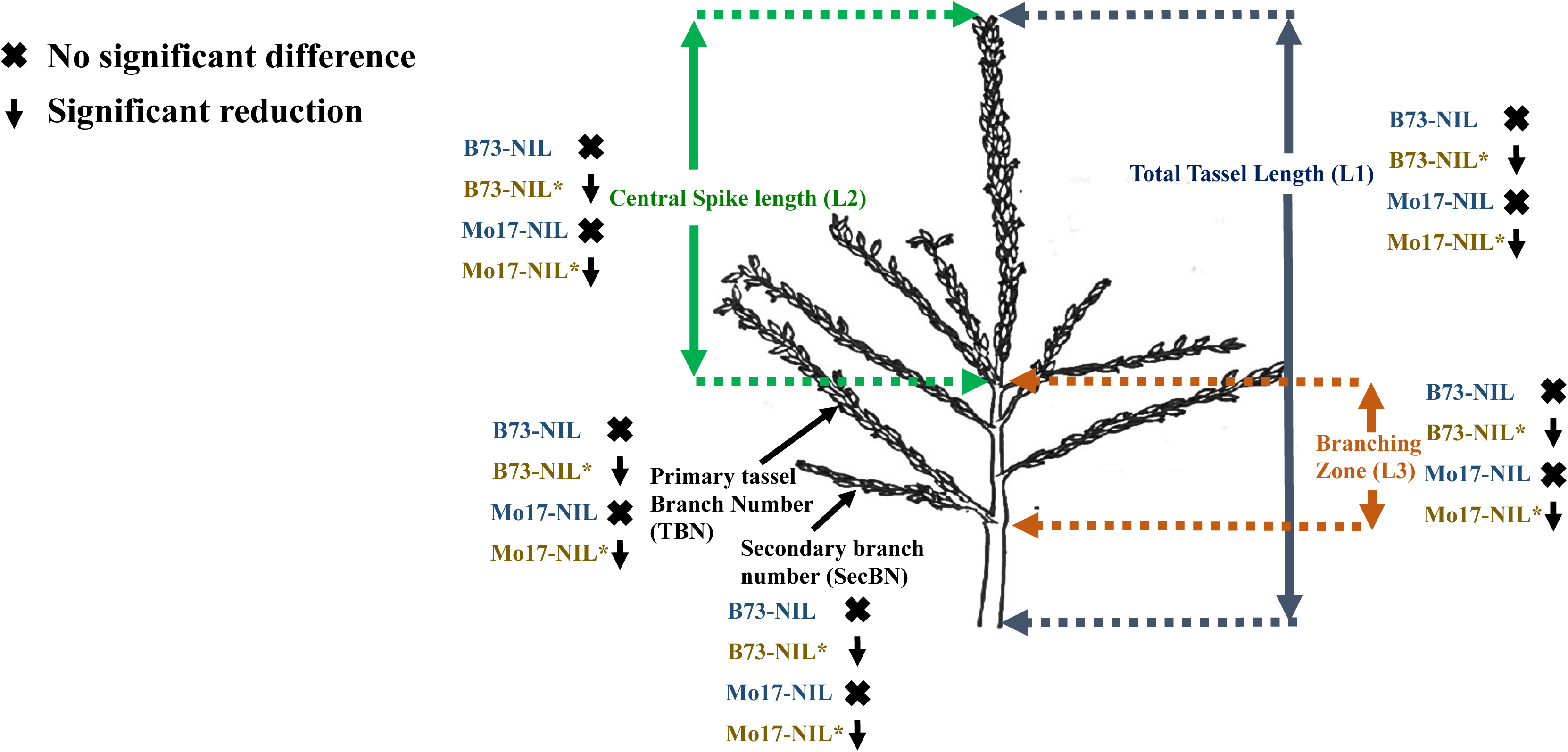
The epistatic effect of *vey1^Mo17^* allele on tassel morphology of *Oy1-N1989* mutants in B73 and B73 x Mo17 isogenic backgrounds. The data presented on the schematic of the maize tassel is derived from mutant siblings of B73 and Mo17-like NILs that were crossed to *Oy1-N1989/+*:B73 tester. Blue colored B73 and Mo17-NILs carry *vey1^B73^* allele, whereas a golden color with asterisk represents NILs with *vey1^Mo17^* introgression. The black cross and downward pointing arrow in a give tassel trait represents no significant reduction and a statistically significant reduction in mutants compared to the wildtype siblings, respectively. Full data is provided in Table S9.

### Tillering is reduced in *Oy1-N1989* mutants and this effect is modulated by *vey1*

Sugar availability is implicated in axillary bud outgrowth (Barbier *et al*. 2015). The levels of sucrose and other reducing sugars are lower in *Oy1-N1989* and further modified by *vey1* (Khangura *et al*. 2019b). We tested if *Oy1-N1989* modulated tiller outgrowth by crossing the mutant to well characterized prolific tillering genetic stocks such as (a) the maize inbred P39 which carries the high tillering allele at *tiller number1* (Zhang *et al*. 2019), (b) the classical dominant tillering mutant *Tillering1*, and (c) a semi-dominant *tb1* in a mixed background. The tillering of the F_1_ hybrids between B73 and all three high tillering lines (sweet corn inbred P39, *Tlr1*, and *tb1*) was suppressed by *Oy1-N1989* allele (**Figure 9** **and Table S10**). To eliminate mixed and hybrid genetic background effects in these crosses, we further tested the influence of *Oy1-N1989* mutation on tillering in *tb1* and *gt1* mutants in isogenic B73 backgrounds. We used B73-like NIL carrying both *vey1* alleles to generate F2 populations that segregated for *Oy1-N1989* along with *tb1* or *gt1* mutants in an isogenic B73 background. The *Oy1-N1989* allele suppressed tillering in both *tb1* and *gt1* mutants. The effect of *Oy1-N1989* on tillering was further modified by *vey1* in the *tb1;Oy1-N1989*/+ and *gt1;Oy1-N1989*/+ double mutants as compared to *tb1* and *gt1* mutant siblings (**Figure 10**). The *tb1;Oy1-N1989*/+ and *gt1;Oy1-N1989*/+ plants that inherited *vey1^B73^* alleles from the NIL had an intermediate number of tillers whereas those with *vey1^Mo17^* alleles had little to no lateral tiller outgrowth. Though *Oy1-N1989/+* suppressed tillering in *gt1* homozygotes, these plants were readily distinguishable from *gt1* heterozygotes and wildtype siblings due to extended blade tissue from the husk sheath (**Figure 10b**) that was described previously (Whipple *et al*. 2011). Thus, the enhancement and suppression of tiller outgrowth in *tb1* and *gt1* mutants by the *Oy1-N1989* and *vey1* alleles displayed the additive and linear relationship previously observed for chlorophyll accumulation and net CO_2_ fixation. This is consistent with sugar availability controlling tiller outgrowth in maize (Mason *et al*. 2014; Barbier *et al*. 2015). The possibility that these changes in axillary branch growth in the combinations of *tin1;Oy1-N1989*/+, *tb1;Oy1-N1989*/+ and *gt1;Oy1-N1989*/+ are due to change in photosynthetic capacity, suggest strongly that future investigations seeking identification of relationships between the fundamentals of energy balance sensing and lateral organ growth control are needed.

**Figure 9.**
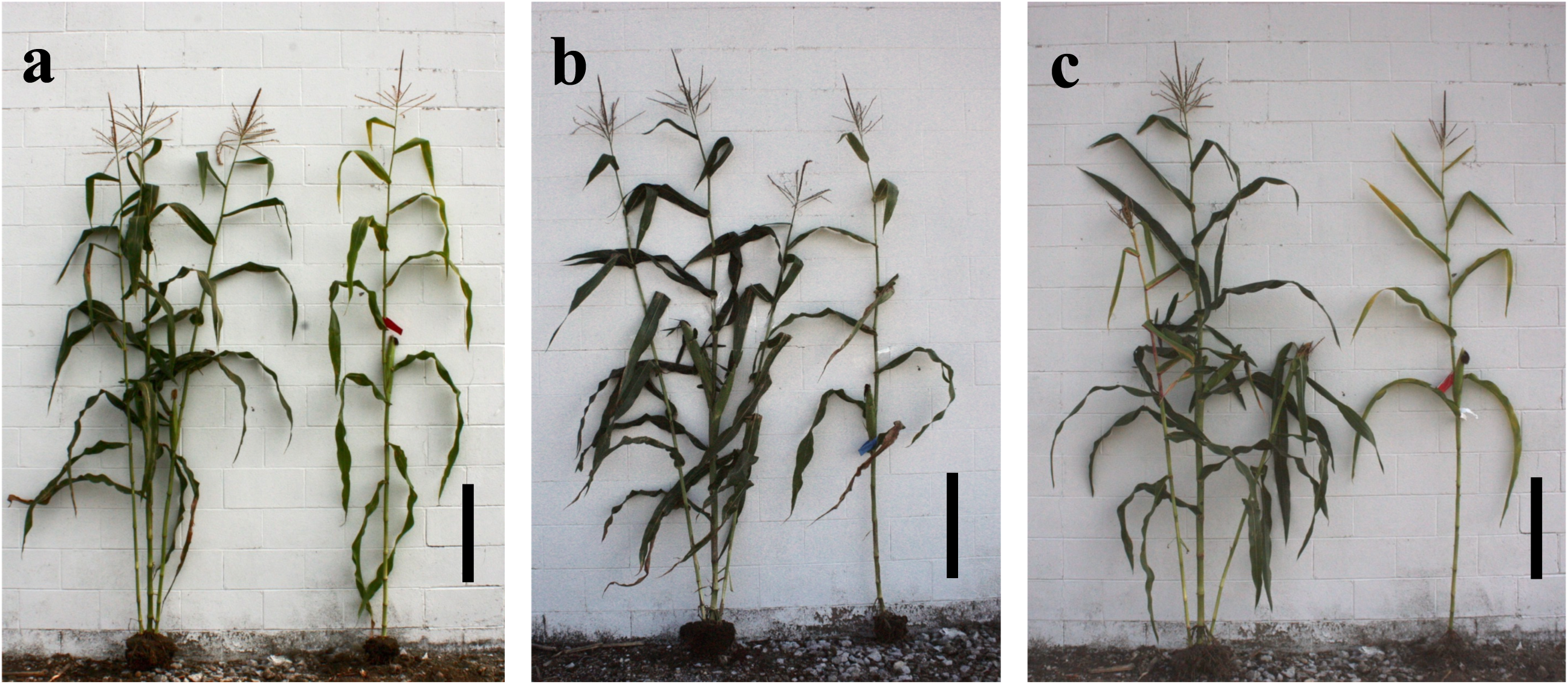
The *Oy1-N1989* allele suppresses tillering in maize. Representative plants from a cross between (a) *Tlr1/Tlr1*;*+/+* x *+/+*;*Oy1-N1989/+*:B73, photographed left to right: *Tlr1/+*;*+/+* and *Tlr1/+*;*Oy1-N1989/+*. (b) *+/+*:P39 x *Oy1-N1989/+*:B73, left to right: *+/+*:P39/B73 and *Oy1-N1989/+*:P39/B73. (c) *tb1-ref/tb1-ref*;*+/+* x *+/+*;*Oy1-N1989/+*:B73, left to right: *tb1-ref/+*;*+/+* and *tb1-ref/+*;*Oy1-N1989/+*. Scale bar in each figure is 50 cm.

**Figure 10.**
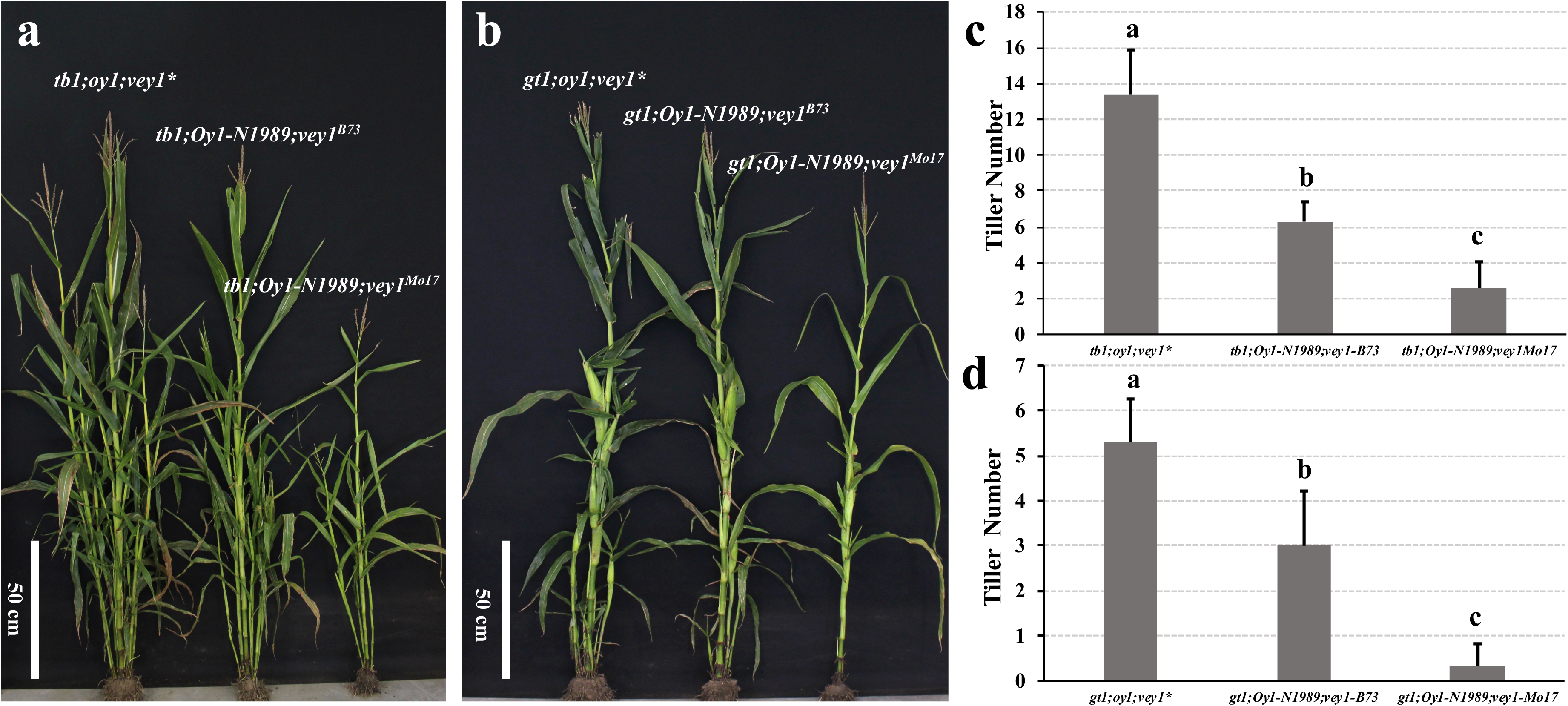
The *vey1* alleles show an additive effect on tillering in the double mutants of the *Oy1-N1989* allele with both *tb1*and *gt1* mutants. Representative (a) *tb1*, and (b) *gt1* mutant plants with different allelic combinations at *vey1* locus forming three distinct genotypic classes. Tiller counts of these three genotypic classes in both (c) *tb1* and (d) *gt1* mutant backgrounds. The connecting letter report in panel c and d indicate statistical significance determined using the student’s t-test at p<0.01. *vey1*\* denotes combination of *vey1^B73^* and *vey1^Mo17^* genotypes.

## Discussion

We demonstrate here that modulation of chlorophyll accumulation in *Oy1-N1989* mutants by the standing variation in the maize germplasm at *vey1* varies metabolism sufficient to alter plant growth and development. Our previous study using these genetic materials showed that cis-acting expression polymorphism at *oy1* locus encoded by alleles of *vey1* can influence the transcript abundance of wildtype OY1. As a result, relative increase in wildtype OY1 transcripts by suppressing *vey1* alleles was consistent with higher chlorophyll accumulation in *Oy1-N1989*/+ mutants, and vice-versa (Khangura *et al*. 2019a). We further demonstrated that the lower chlorophyll levels in *Oy1-N1989* mutants conditioned by *vey1* alleles resulted in lower photosynthetic rate, non-structural carbohydrate synthesis, and delayed reproductive maturity (Khangura *et al*. 2019b). In this study, we demonstrate that chlorophyll metabolism, and the variation affected by the *vey1* locus interactions with the *Oy1-N1989* mutant allele, sufficiently disrupts metabolism to alter stem thickness, plant height, tiller production, and tassel branching. In all traits but plant height, *Oy1-N1989* mutants had lower trait expression than the wildtype siblings, consistent with a loss of growth due to decreased photosynthetic output. The performance of the mutant plants was further reduced for these traits by the enhancing *vey1^Mo17^* allele. So far, all phenotypes except plant height show a linear change in trait value with suppressing (e.g. B73) and enhancing (e.g. Mo17) alleles at *vey1* (**Figures 2, 4, and 5**). This relationship is not a genetic correlation between alleles and phenotypes. As expected for the traits to be determined by the effect of alleles on chlorophyll biosynthesis, the variation in chlorophyll levels between individuals carrying the same *vey1* genotype were also correlated with consistent changes in plant phenotypes for all traits other than plant height (**Figures S3-S9**).

The effect of *vey1* alleles on mutant plant height does not follow a linear relationship and cannot be explained by continuous loss of chlorophyll creating a continuum of sugar levels which support differing plant growth. The *Oy1-N1989/+* mutants inheriting a *vey1^B73^* allele had a mild decrease in chlorophyll and lower net CO_2_ assimilation (Khangura *et al*. 2019a, 2019b) and yet were taller than their wildtype siblings. The *Oy1-N1989/+* mutants with the *vey1^Mo17^* allele that have further reduction in chlorophyll and net CO_2_ assimilation were shorter than their wildtype siblings (**Figures 2, 4, and 5; Table S4**). These results contrast to those from *Arabidopsis*, where a 20% decrease in chlorophyll lead to a small but significant decrease in hypocotyl elongation under the shade and a further decrease lead to greater reduction in hypocotyl lengths (Ortiz-Alcaide *et al*. 2019). In our experiments neither the ∼80% nor 90% reduction in chlorophyll of the *Oy1-N1989/+* mutants with the *vey1^B73^* and *vey1^Mo17^* alleles (from the CCMI measurements in Khangura et al., 2019b) reduced nor increased mesocotyl elongation under continuous white light, darkness, red, or far-red light. (**Figure 7**). This may simply relate to the amount of carbohydrates stored in the maize seed available for mesocotyl elongation. The opposing effects of *vey1* alleles on the influence of *Oy1-N1989* on plant height strongly argues for additional experiments to distinguish whether a biochemical signal related to tetrapyrrole metabolism or an indirect effect of chlorophyll loss, unrelated to CO_2_ assimilation, lead to the changes in plant height.

In addition to *vey1*, multiple QTLs were detected for stalk width and plant height measurements in both mutant and wildtype siblings. Putative genes underlying these QTLs discussed above could explain the genetics of these traits. One of the QTL for total plant height variation in the wildtype siblings in the Syn10 F_1_ population on chromosome 3 contained *zmmads69* (**Table S8**). QTL for plant height and flowering time in the IBM and Syn10 inbred populations were previously mapped to a similar position (Liu *et al*. 2015). Recently, *zmmads69* was characterized as a positive regulator of flowering time and was proposed as a component of the *ZmRap2.7-ZCN8* regulatory network that allowed latitudinal adaptation of maize (Liang *et al*. 2019). Overexpression of *zmmads69* lead to early flowering time and also reduced plant and ear height (Liang *et al*. 2019) and a strong cis-eQTL for *zmmads69* (LOD=44.17; R^2^=0.76) was detected in the IBM inbred lines, with the Mo17 allele resulting in greater transcript abundance, reduced plant height, and earlier maturity. We demonstrate here that the Mo17 allele at this locus is associated with a QTL that decreased plant height in the wildtype siblings of our Syn10 F_1_ population (**Table S8**) and, when considered without a multiple test correction, decreased days to anthesis.

Since we only measured tillering and tassel architecture in NIL crosses, it is a formal possibility that a modifier of plant branching is linked to *oy1*. For example, a tillering modifier that is common to both the *gt1* and *tb1* responses may be linked to *oy1*. However, the changes in the developmental phenotypes were contingent on *Oy1-N1989/+* and it retained the epistatic interaction with *vey1* that we have found for all traits (**Figure 10**). This makes it very unlikely that a linked QTL differing between B73 and Mo17 underlie this variation in tillering. In the case of tassel branching, our results are an independent validation of a previous report of reduced tassel branching by a loss-of-function allele at *oy1* (Brewbaker 2015). In all cases, lateral branch numbers were reduced by loss of chlorophyll regardless of the modifying allele at *vey1* (**Figure 8**).

The maize gene, *tiller number1* (*tin1*) encodes a C2H2-zinc-finger transcription factor which is a dominant factor responsible for increased tiller growth in the Purdue sweetcorn line P51 (Zhang *et al*. 2019). TIN1 is upstream of GT1, but acts independently of TB1 (Kebrom and Brutnell 2015; Zhang *et al*. 2019). The high tillering Purdue Sweetcorn line P39, encodes the same *tin1* allele as P51 (Zhang *et al*. 2019) possibly due to their common derivation from ‘Golden Bantam’ (www.ars-grin.gov). The *Oy1-N1989* allele suppressed tillering conditioned by *tin1* variant in P39 (**Figure 9**). The suppression of tillering in *gt1* mutants in *Oy1-N1989/+* backgrounds (**Figure 10**) is consistent with *tin1* acting upstream of *gt1* (Zhang *et al*. 2019) to regulate tiller bud out growth. *Oy1-N1989* also suppressed *tb1* despite the fact that tb1 acts independently of *tin1,* suggests that loss of chlorophyll regulates branching upstream of the branch point between *tb1* and *tin1-gt1* (**Figures 9** and **10**; **Table S10**). In isogenic double mutants of *Oy1-N1989* allele with *gt1* and *tb1*, *vey1* locus that have been previously shown to cause gradual decline in photosynthetic capacity and sugar accumulation of *Oy1-N1989* mutants (Khangura *et al*. 2019b) also influenced tiller number in a *vey1*-dependent manner (**Figure 10**). A study analyzing young axillary buds of *tb1* and *gt1* mutants in maize found increased sugar biosynthesis and metabolism to be critical for breaking bud dormancy and that *tb1* bound to the promoters of genes encoding enzymes important for energy sensing, such as enzymes involved in trehalose-6-phosphate metabolism (Dong *et al*. 2019). Analysis of transcript abundance in axillary buds of *tin1* variants found photosynthesis to be the most enriched gene ontology term and chlorophyll biosynthesis to be highly upregulated in non-dormant tiller buds (Zhang *et al*. 2019). This may be tempting to dismiss, as the initiation of photosynthesis and respiration must be a consequence of bud outgrowth. However, it may be that sugar production at the branches is a critical determinant of branch growth. Based on our data that chlorophyll limitation suppresses tillering by *tb1*, *gt1*, and *tin1* alleles, we predict that these genes regulate the process of bud outgrowth via stem sucrose concentrations and that these transcription factors act downstream to modify bud growth. We expect genes important for the patterning and placement of axillary meristems to act upstream of the sensor of carbon status.

Flowering time and plant height displayed a confusing relationship that was not resolved by the idea that later flowering plants were taller. Indeed, as expected based on the correlations with CCM and *vey1* but in defiance of a simple explanation, as mutant plants accumulated more chlorophyll they flowered earlier and were taller in IBM regardless of the *vey1* genotype (**Figure S8**). In the Syn10 F_1_ population, the plants with *vey1^Mo17^* allele behaved in this manner, but the *vey1^B73^* plants went in the opposite direction with increasing height and increasing flowering time co-occurring (**Figure S9**). Thus, a secondary relationship between plant height and the flowering time delay does not explain the non-linear change in plant height with respect to chlorophyll levels. Although the plant height in *Oy1-N1989* and *vey1* segregating materials is extremely reproducible across range of populations (two RIL populations and NIL) and across multiple years, the mechanism that explains the complicated relationship between the *Oy1-N1989* allele, its modifier *vey1* and CCM values to plant height and the paradoxical effect of this epistasis on different morphological traits is elusive to us at this time.

There is some separation between the CCM values that are produced by *vey1^B73^-Oy1-N1989* and those produced by *vey1^Mo17^-Oy1-N1989.* It may be that a “goldilocks” value for CCM that maximizes plant height sits between the values we have measured. But if so, *Oy1-N1989/+* mutants suppressed by a *vey1^B73^* allele should get taller as CCM decreases. Instead, the *vey1^B73^* plants show no significant correlations between CCM and plant height with both positive and negative slope of the regression line consistent with the asymptotic relationship between CCM and plant height. If there is a break-even point where decreasing chlorophyll contents increased plant height, it occurs between the CCM values of the two *vey1* alleles. Thus far, the relationship between *vey1* and plant height and the unusual effect of *vey1^B73^* on plant height denies any simple explanation. Though not measured by us, a shorter thinner plant should produce less biomass and utilize less carbohydrate. The plant volume which is a function of stem thickness and plant height is built from excess CO_2_ assimilation. As a result, it may be that the *vey1* suppressed *Oy1-N1989*/+ mutants are taller because of the addition of available carbohydrate to thinner stems. Direct measurements of biomass might better estimate this and would also open up the possibility to explore other aspects such as water and nitrogen use efficiency.

## Acknowledgements

We would like to thank Dr. Sujith Puthiyaveetil and Dr. Iskander Ibrahim on their scientific insight and help with conducting the light quality experiment. Dr. Brenda Owens is gratefully acknowledged for her botanical illustration of a maize tassel.

## References

1. Austin, D. F., M. Lee, and L. R. Veldboom, 2001 Genetic mapping in maize with hybrid progeny across testers and generations: Plant height and flowering. Theor. Appl. Genet. 102: 163–176.

2. Barbier, F. F., J. E. Lunn, and C. a Beveridge, 2015 Ready, steady, go! A sugar hit starts the race to shoot branching. Curr. Opin. Plant Biol. 25: 39–45.

3. Beavis, W. D., 1994 The power and deceit of QTL experiments: lessons from comparative QTL studies, pp. 252–268 in 49th Annual Corn and Sorghum Research Conference, American Seed Trade Association,.

4. Beavis, W. D., D. Grant, M. Albertsen, and R. Fincher, 1991 Quantitative trait loci for plant height in four maize populations and their associations with qualitative genetic loci. Theor Appl Genet 83: 141–145.

5. Beavis, W. D., O. S. Smith, D. Grant, and R. Fincher, 1994 Identification of quantitative trait loci using a small sample of topcrossed and F4 progeny from maize. Crop Sci. 34: 882–896.

6. Brewbaker, J. L., 2015 Diversity and Genetics of Tassel Branch Numbers in Maize. Crop Sci. 55:65.

7. Broman, K. W., H. Wu, S. Sen, and G. a. Churchill, 2003 R/qtl: QTL mapping in experimental crosses. Bioinformatics 19: 889–890.

8. Chaikam, V., A. Negeri, R. Dhawan, B. Puchaka, J. Ji et al., 2011 Use of Mutant-Assisted Gene Identification and Characterization (MAGIC) to identify novel genetic loci that modify the maize hypersensitive response. Theor. Appl. Genet. 123: 985–97.

9. Chen, M., J. Chory, and C. Fankhauser, 2004 Light Signal Transduction in Higher Plants. Annu. Rev. Genet. 38: 87–117.

10. Churchill, G. A., and R. W. Doerge, 1994 Empirical threshold values for quantitative trait mapping. Genetics 138: 963–71.

11. Dong, Z., Y. Xiao, R. Govindarajulu, R. Feil, M. L. Siddoway et al., 2019 The regulatory landscape of a core maize domestication module controlling bud dormancy and growth repression. Nat. Commun. 10:.

12. Eichten, S. R., J. M. Foerster, N. de Leon, Y. Kai, C.-T. Yeh et al., 2011 B73-Mo17 near-isogenic lines demonstrate dispersed structural variation in maize. Plant Physiol. 156: 1679–90.

13. Eyster, W., 1933 Linkage studies in maize. Am. Nat. 67: 75.

14. Forestan, C., S. Farinati, and S. Varotto, 2012 The maize PIN gene family of auxin transporters. Front. Plant Sci. 3: 1–23.

15. Frankenberg, N., and J. C. Lagarias, 2012 Biosynthesis and Biological Functions of Bilins. Elsevier Inc.

16. Gibson, L. C. D., R. D. Willows, C. G. Kannangara, D. von Wettstein, and C. N. Hunter, 1995 Magnesium-protoporphyrin chelatase of Rhodobacter sphaeroides: reconstitution of activity by combining the products of the bchH, -I, and -D genes expressed in Escherichia coli. Proc. Natl. Acad. Sci. U. S. A. 92: 1941–1944.

17. Govindjee, D. Shevela, and L. O. Björn, 2017 Evolution of the Z-scheme of photosynthesis: a perspective. Photosynth. Res. 133: 5–15.

18. Guo, Y., H. Wu, X. Li, Q. Li, X. Zhao et al., 2017 Identification and expression of GRAS family genes in maize (Zea mays L.). PLoS One 12: 1–17.

19. Hirsch, C. N., J. M. Foerster, J. M. Johnson, R. S. Sekhon, G. Muttoni et al., 2014 Insights into the maize pan-genome and pan-transcriptome. Plant Cell 26: 121–135.

20. Kebrom, T. H., and T. P. Brutnell, 2015 Tillering in the sugary1 sweetcorn inbred is maintained by overriding the teosinte branched1 repressive signal. Plant Signal. Behav. 10: 1–4.

21. Khangura, R. S., S. Marla, B. P. Venkata, N. J. Heller, G. S. Johal et al., 2019 A Very Oil Yellow1 Modifier of the Oil Yellow1-N1989 Allele Uncovers a Cryptic Phenotypic Impact of Cis-regulatory Variation in Maize. G3 9: 375–390.

22. Khangura, R. S., B. P. Venkata, S. R. Marla, M. V Mickelbart, S. Dhungana et al., 2019a Interaction between induced and natural variation at oil yellow1 delays flowering in maize. bioRxiv.

23. Khangura, R. S., B. P. Venkata, S. R. Marla, M. V Mickelbart, S. Dhungana et al., 2019b Interaction between induced and natural variation at oil yellow1 delays reproductive maturity in maize. G3 1–48.

24. Kremling, K. A. G., S.-Y. Chen, M.-H. Su, N. K. Lepak, M. C. Romay et al., 2018 Dysregulation of expression correlates with rare-allele burden and fitness loss in maize. Nature 555: 520– 523.

25. Lee, Y., and H. Kende, 2001 Expression of B-Expansins Is Correlated with Internodal Elongation in Deepwater Rice. Plant Physiol. 127: 645–654.

26. Lemoine, R., S. La Camera, R. Atanassova, F. Dédaldéchamp, T. Allario et al., 2013 Source-to-sink transport of sugar and regulation by environmental factors. Front. Plant Sci. 4: 1–21.

27. Liang, Y., Q. Liu, X. Wang, C. Huang, G. Xu et al., 2019 ZmMADS69 functions as a flowering activator through the ZmRap2.7-ZCN8 regulatory module and contributes to maize flowering time adaptation. New Phytol. 221: 2335–2347.

28. Li, L., K. Petsch, R. Shimizu, S. Liu, W. W. Xu et al., 2018 Correction: Mendelian and Non-Mendelian Regulation of Gene Expression in Maize. PLoS Genet. 14: e1007234.

29. Li, L., K. Petsch, R. Shimizu, S. Liu, W. W. Xu et al., 2013 Mendelian and non-Mendelian regulation of gene expression in maize. PLoS Genet. 9: e1003202.

30. Lin, Hying, Q. Liu, X. Li, J. Yang, S. Liu et al., 2017 Substantial contribution of genetic variation in the expression of transcription factors to phenotypic variation revealed by eRD-GWAS. Genome Biol. 18: 1–14.

31. Liu, H., Y. Niu, P. J. Gonzalez-Portilla, H. Zhou, L. Wang et al., 2015 An ultra-high-density map as a community resource for discerning the genetic basis of quantitative traits in maize. BMC Genomics 16: 1078.

32. Ma, Z., X. Hu, W. Cai, W. Huang, X. Zhou et al., 2014 Arabidopsis miR171-Targeted Scarecrow-Like Proteins Bind to GT cis-Elements and Mediate Gibberellin-Regulated Chlorophyll Biosynthesis under Light Conditions. PLoS Genet. 10: 20–21.

33. Macadlo, L. A., I. M. Ibrahim, and S. Puthiyaveetil, 2019 Sigma factor 1 in chloroplast gene transcription and photosynthetic light acclimation. J. Exp. Bot. 1–10.

34. Mason, M. G., J. J. Ross, B. A. Babst, B. N. Wienclaw, and C. A. Beveridge, 2014 Sugar demand, not auxin, is the initial regulator of apical dominance. Proc. Natl. Acad. Sci. 111: 6092–6097.

35. Minow, M. A. A., L. M. Ávila, K. Turner, E. Ponzoni, I. Mascheretti et al., 2018 Distinct gene networks modulate floral induction of autonomous maize and photoperiod-dependent teosinte. J. Exp. Bot. 69: 2937–2952.

36. Ortiz-Alcaide, M., E. Llamas, A. Gomez-Cadenas, A. Nagatani, J. F. Martínez-García et al., 2019 Chloroplasts modulate elongation responses to canopy shade by retrograde pathways involving hy5 and abscisic acid. Plant Cell 31: 384–398.

37. Pierik, R., and M. De Wit, 2014 Shade avoidance: Phytochrome signalling and other aboveground neighbour detection cues. J. Exp. Bot. 65: 2815–2824.

38. Puthiyaveetil, S., 2008 Two-component signalling systems of chloroplasts: function, distribution and evolution: Queen Mary University of London, Ph.D. Dissertation p.

39. Rabinowitch, E. I., and Govindjee, 1965 The Role of Chlorophyll in Photosynthesis. Sci. Am. 213: 74–83.

40. Roig-Villanova, I., and J. F. Martínez-García, 2016 Plant responses to vegetation proximity: A whole life avoiding shade. Front. Plant Sci. 7: 1–10.

41. Sawers, R. J. H., P. J. Linley, P. R. Farmer, N. P. Hanley, D. E. Costich et al., 2002 elongated mesocotyl1, a Phytochrome-Deficient Mutant of Maize. Plant Physiol. 130: 155–163.

42. Sawers, R. J. H., P. J. Linley, J. F. Gutierrez-Marcos, T. Delli-Bovi, P.R. Farmer et al., 2004 The Elm1 (ZmHy2) gene of maize encodes a phytochromobilin synthase. Plant Physiol. 136: 2771–2781.

43. Sawers, R. J. H., J. Viney, P. R. Farmer, R. R. Bussey, G. Olsefski et al., 2006 The maize Oil yellow1 (Oy1) gene encodes the I subunit of magnesium chelatase. Plant Mol. Biol. 60: 95– 106.

44. Springer, P. S., and J. L. Bennetzen, 1995 Tlr1 may be allelic to tb1. Maize Genet. Coop. Newsl. 69: 136.

45. Vernier, P., 1631 La construction, l’usage, et les propriétés du quadrant nouveau de mathématique. Brussels.

46. Walker, B. J., D. T. Drewry, R. A. Slattery, A. VanLoocke, Y. B. Cho et al., 2017 Chlorophyll can be reduced in crop canopies with little penalty to photosynthesis. Plant Physiol. 176: 1215– 1232.

47. Walker, C. J., and J. D. Weinstein, 1991 In vitro assay of the chlorophyll biosynthetic enzyme Mg-chelatase: resolution of the activity into soluble and membrane-bound fractions. Proc. Natl. Acad. Sci. U. S. A. 88: 5789–5793.

48. Wang, R., X. Liu, S. Liang, Q. Ge, Y. Li et al., 2015 A subgroup of MATE transporter genes regulates hypocotyl cell elongation in Arabidopsis. J. Exp. Bot. 66: 6327–6343.

49. Wang, B., H. Liu, Z. Liu, X. Dong, J. Guo et al., 2018 Identification of minor effect QTLs for plant architecture related traits using super high density genotyping and large recombinant inbred population in maize (Zea mays). BMC Plant Biol. 18: 1–12.

50. Wang, Y., J. Yao, Z. Zhang, and Y. Zheng, 2006 The comparative analysis based on maize integrated QTL map and meta-analysis of plant height QTLs. Chinese Sci. Bull. 51: 2219– 2230.

51. Waters, A. J., I. Makarevitch, J. Noshay, L. T. Burghardt, C. N. Hirsch et al., 2017 Natural variation for gene expression responses to abiotic stress in maize. Plant J. 89: 706–717.

52. Whipple, C. J., T. H. Kebrom, A. L. Weber, F. Yang, D. Hall et al., 2011 Grassy tillers1 promotes apical dominance in maize and responds to Shade signals in the grasses. Proc. Natl. Acad. Sci. U. S. A. 108: 506–512.

53. Winkler, R. G., and T. Helentjaris, 1995 The maize Dwarf3 gene encodes a cytochrome P450-mediated early step in gibberellin biosynthesis. Plant Cell 7: 1307–1317.

54. Yu, S. M., S. F. Lo, and T. H. D. Ho, 2015 Source-Sink Communication: Regulated by Hormone, Nutrient, and Stress Cross-Signaling. Trends Plant Sci. 20: 844–857.

55. Zhang, X., Z. Lin, J. Wang, H. Liu, L. Zhou et al., 2019 The tin1 gene retains the function of promoting tillering in maize. Nat. Commun. 10: 1–13.

56. Zhang, H., H. Zhu, Y. Pan, Y. Yu, S. Luan et al., 2014 A DTX/MATE-type transporter facilitates abscisic acid efflux and modulates ABA sensitivity and drought tolerance in Arabidopsis. Mol. Plant 7: 1522–1532.

