## Supplemental Tables S4-S10 for "Variation in maize chlorophyll biosynthesis alters plant architecture"

Table S4. Effect of *Oy1-N1989* and genetic background on chlorophyll index, plant height traits, and stalk width.

| Background | Genotype | CCMI | CCMII | SW | FlHT | EaHT | Ea2Fl |
| --- | --- | --- | --- | --- | --- | --- | --- |
| B73 | *+/+* | 27.5±2.4^a^ | 64.6±8.0^a^ | 22.3±1.4^a^ | 200.5±5.4^a^ | 95.2±5.5^a^ | 105.3±3.8^a^ |
| Mo17 x B73 | *+/+* | 31.2±6.1^a^ | 78.2±5.8^b^ | 28.1±1.1^b^ | 228.8±8.5^b^ | 107.5±3.5^b^ | 121.3±7.2^b^ |
| B73 | *Oy1-1989/+* | 4.7±1.2^b^ | 22.2±3.1^c^ | 20.5±1.8^ac^ | 211.3±6.8^c^ | 107.1±2.3^b^ | 104.1±7.9^a^ |
| Mo17 x B73 | *Oy1-1989/+* | 2.0±0.4^c^ | 5.7±1.1^d^ | 18.8±1.5^c^ | 232.3±4.9^b^ | 100.6±5.3^a^ | 131.7±6.4^c^ |

Each value is an average± standard deviation of plants derived from five replications planted in randomized complete block design. Connecting letter report indicates statistical significance determined using student’s t-test with p<0.05.

Table S5. Pairwise correlation calculations of traits measured in IBM x *Oy1-N1989/+*:B73 F_1_ population

|  | **MT_CCMII** | **WT_CCMII** | **MT_FlHT** | **WT_FlHT** | **Diff_FlHT** | **MT_SW** | **WT_SW** | **Diff_SW** | **MT_DTA** | **WT_DTA** |
| --- | --- | --- | --- | --- | --- | --- | --- | --- | --- | --- |
| **MT_CCMII** |  | 0.19 | 0.35 | 0.03 | -0.60 | 0.64 | -0.03 | -0.76 | -0.30 | 0.13 |
| **WT_CCMII** | 0.005 |  | 0.32 | 0.35 | -0.06 | 0.29 | 0.46 | 0.03 | 0.37 | 0.36 |
| **MT_FlHT** | <.0001 | <.0001 |  | 0.84 | -0.51 | 0.63 | 0.57 | -0.27 | 0.49 | 0.73 |
| **WT_FlHT** | 0.67 | <.0001 | <.0001 |  | 0.04 | 0.40 | 0.70 | 0.10 | 0.72 | 0.82 |
| **Diff_FlHT** | <.0001 | 0.38 | <.0001 | 0.55 |  | -0.53 | 0.03 | 0.64 | 0.21 | -0.07 |
| **MT_SW** | <.0001 | <.0001 | <.0001 | <.0001 | <.0001 |  | 0.52 | -0.72 | 0.14 | 0.51 |
| **WT_SW** | 0.61 | <.0001 | <.0001 | <.0001 | 0.61 | <.0001 |  | 0.22 | 0.76 | 0.76 |
| **Diff_SW** | <.0001 | 0.62 | <.0001 | 0.15 | <.0001 | <.0001 | 0.0008 |  | 0.44 | 0.02 |
| **MT_DTA** | <.0001 | <.0001 | <.0001 | <.0001 | 0.002 | 0.04 | <.0001 | <.0001 |  | 0.83 |
| **WT_DTA** | 0.06 | <.0001 | <.0001 | <.0001 | 0.32 | <.0001 | <.0001 | 0.75 | <.0001 |  |

Values above the diagonal are Pearson’s correlation coefficients, and values below the diagonal are P-values for pairwise comparisons.

Table S6. Pairwise correlation calculations of traits measured in Syn10 x *Oy1-N1989/+*:B73 F_1_ population

|  | **WT_CCMII** | **MT_CCMII** | **WT_DTA** | **MT_DTA** | **WT_SW** | **MT_SW** | **Diff_SW** | **WT_FlHT** | **MT_FlHT** | **Diff_FlHT** | **WT_EaHT** | **MT_EaHT** | **Diff_EaHT** |
| --- | --- | --- | --- | --- | --- | --- | --- | --- | --- | --- | --- | --- | --- |
| **WT_CCMII** |  | 0.01 | -0.05 | 0.03 | 0.05 | -0.01 | 0.03 | -0.15 | -0.01 | -0.11 | -0.15 | 0.00 | -0.13 |
| **MT_CCMII** | 0.93 |  | -0.03 | -0.90 | -0.21 | 0.86 | -0.88 | 0.04 | 0.55 | -0.65 | 0.06 | 0.42 | -0.51 |
| **WT_DTA** | 0.44 | 0.65 |  | 0.20 | 0.04 | -0.04 | 0.05 | 0.29 | 0.17 | 0.04 | 0.30 | 0.21 | 0.00 |
| **MT_DTA** | 0.65 | <.0001 | 0.002 |  | 0.23 | -0.85 | 0.88 | 0.04 | -0.48 | 0.63 | 0 | -0.33 | 0.45 |
| **WT_SW** | 0.47 | 0.001 | 0.52 | 0.0002 |  | 0.02 | 0.37 | 0.27 | 0.07 | 0.15 | 0.15 | 0.04 | 0.09 |
| **MT_SW** | 0.88 | <.0001 | 0.52 | <.0001 | 0.77 |  | -0.92 | 0.11 | 0.61 | -0.67 | 0.14 | 0.48 | -0.52 |
| **Diff_SW** | 0.68 | <.0001 | 0.44 | <.0001 | <.0001 | <.0001 |  | 0.01 | -0.55 | 0.69 | -0.07 | -0.43 | 0.51 |
| **WT_FlHT** | 0.02 | 0.57 | <.0001 | 0.54 | <.0001 | 0.09 | 0.93 |  | 0.60 | 0.12 | 0.83 | 0.60 | -0.03 |
| **MT_FlHT** | 0.85 | <.0001 | 0.01 | <.0001 | 0.30 | <.0001 | <.0001 | <.0001 |  | -0.72 | 0.54 | 0.87 | -0.66 |
| **Diff_FlHT** | 0.08 | <.0001 | 0.54 | <.0001 | 0.02 | <.0001 | <.0001 | 0.05 | <.0001 |  | 0.05 | -0.55 | 0.80 |
| **WT_EaHT** | 0.02 | 0.33 | <.0001 | 1.00 | 0.02 | 0.03 | 0.26 | <.0001 | <.0001 | 0.43 |  | 0.67 | 0.05 |
| **MT_EaHT** | 0.97 | <.0001 | 0.001 | <.0001 | 0.55 | <.0001 | <.0001 | <.0001 | <.0001 | <.0001 | <.0001 |  | -0.71 |
| **Diff_EaHT** | 0.03 | <.0001 | 0.97 | <.0001 | 0.16 | <.0001 | <.0001 | 0.66 | <.0001 | <.0001 | 0.47 | <.0001 |  |

Values above the diagonal are Pearson’s correlation coefficients, and values below the diagonal are P-values for pairwise comparisons.

Table S7. The QTL detected in the hybrid IBM x *Oy1-N1989*/+:B73 population

|  |  |  |  |  |  |  |  |  |  | Mean±SE^g^ | |
| --- | --- | --- | --- | --- | --- | --- | --- | --- | --- | --- | --- |
| Trait Identifier | Chr^a^ | Position^b^ | LOD^c^ | Left Marker | Right Marker | Interval (cM)^d^ | Interval  (Mbp)^e^ | Candidate | PVE^f^ | B73 | Mo17 |
| WT_FlHT | 9 | 257 | 7.62 | bnl5.04 | umc1267 | 238-272 | 106.0-106.3 |  | 14.3 | 194.83±1.28 | 205.90±1.31 |
| MT_FlHT | 9 | 298 | 4.06 | bnlg1012 | ufg66 | 273-302 | 118.0-119.4 | *zmgras45* | 8.3 | 197.06±1.73 | 207.55±1.64 |
|  | 10 | 137 | 7.48 | AI795367 | AY109994 | 125-154 | 8.7-9.4 | *oy1* | 15.1 | 207.98±1.47 | 193.65±1.91 |
| Ratio_FlHT | 10 | 139 | 16.92 | AI795367 | AY109994 | 135-145 | 8.7-9.4 | *oy1* | 28.5 | 1.04±0.01 | 0.97±0.01 |
| WT_SW | 1 | 810 | 6.76 | nfc103a | AY109506 | 802-821 | 254.4-255.9 | *expb4* | 13.4 | 23.31±0.15 | 24.4±0.12 |
| MT_SW | 10 | 144 | 26.91 | phi059 | isu85b | 137-148 | 8.8-9.4 | *oy1* | 43.3 | 22.37±0.19 | 18.56±0.23 |
| Ratio_SW | 10 | 145 | 35.32 | isu85b | rz900c(ahh) | 126-147 | 8.8-9.4 | *oy1* | 52.6 | 0.94±0.01 | 0.76±0.01 |

^a^Chromosome location of a QTL; ^b^Peak position of the QTL; ^c^LOD score at the peak of a given QTL

^d^2-LOD interval of the QTL in terms of genetic position in centiMorgan(cM); ^e^Physical position (in Mbp) of 2-LOD flanking markers from B73 RefGenV4; ^f^Phenotypic variance explained by the QTL at the peak position as estimated by regression and reported as an R^2^ value*100; ^g^Mean and standard error of the trait with B73 and Mo17 genotype at the peak marker of the detected QTL

Table S8. The QTL detected in the hybrid Syn10 x *Oy1-N1989*/+:B73 population

|  |  |  |  |  |  |  |  |  |  | Mean±SE^g^ | |
| --- | --- | --- | --- | --- | --- | --- | --- | --- | --- | --- | --- |
| Trait Identifier | Chr^a^ | Position^b^ | LOD^c^ | Left Marker | Right Marker | Interval (cM)^d^ | Interval (Mbp)^e^ | Candidate | PVE^f^ | B73 | Mo17 |
| WT_SW | 1 | 1645.0 | 6.2 | chr01.2976.5 | chr01.2977.5 | 1641-1653 | 303.1-303.1 |  | 10.77 | 25.7±0.09 | 24.75±0.14 |
|  | 1 | 785.9 | 5.3 | chr01.1877 | chr01.1878.5 | 774-808 | 189.9-190.0 |  | 9.35 | 25.72±0.1 | 24.86±0.13 |
| MT_SW | 10 | 189.4 | 62.3 | chr10.90.5 | chr10.93 | 188-193 | 8.9-9.1 | *oy1* | 68.23 | 21.74±0.13 | 15.93±0.2 |
| Ratio_SW | 9 | 279.5 | 3.9 | chr09.159.5 | chr09.161 | 261-877 | 15.6-15.8 |  | 6.90 | 0.81±0.01 | 0.75±0.01 |
|  | 10 | 190.0 | 69.1 | chr10.93 | chr10.94.5 | 188-193 | 9.1-9.3 | *oy1* | 70.89 | 0.86±0 | 0.61±0 |
| WT_FlHT | 1 | 538.2 | 3.9 | chr01.519.5 | chr01.522 | 516-553 | 52.2-52.5 |  | 6.86 | 218.4±1.04 | 226.9±1.67 |
|  | 1 | 622.0 | 4.6 | chr01.808 | chr01.809.5 | 612-628 | 82.3-82.5 | MATE | 8.11 | 217.62±1.1 | 226.21±1.45 |
|  | 1 | 779.2 | 5.1 | chr01.1843 | chr01.1860 | 751-832 | 186.2-187.3 |  | 8.90 | 224.45±1.14 | 215.65±1.35 |
|  | 3 | 565.0 | 4.0 | chr03.1591 | chr03.1592.5 | 554-578 | 160.6-160.7 | *mads69* | 7.11 | 225.67±1.43 | 217.74±1.12 |
|  | 9 | 466.0 | 4.4 | chr09.1039.5 | chr09.1050.5 | 458-475 | 106.8-107.9 |  | 7.90 | 217.29±1.16 | 225.45±1.35 |
| MT_FlHT | 1 | 622.0 | 3.8 | chr01.808 | chr01.809.5 | 612-638 | 82.3-82.5 | MATE | 6.67 | 226.5±1.58 | 237.6±2.08 |
|  | 9 | 57.0 | 4.1 | chr09.36.5 | chr09.37.5 | 41-68 | 3.2-3.2 | *pin1a* | 7.49 | 225.77±1.66 | 237.03±1.92 |
|  | 10 | 191.0 | 16.7 | chr10.94.5 | chr10.95.5 | 187-195 | 9.3-9.5 | *oy1* | 26.45 | 237.51±1.33 | 214.28±2.05 |
| Ratio_FlHT | 10 | 189.0 | 24.4 | chr10.90.5 | chr10.93 | 174-194 | 8.9-9.1 | *oy1* | 35.71 | 1.07±0 | 0.97±0 |
| WT_EaHT | 9 | 465.0 | 7.4 | chr09.1039.5 | chr09.1050.5 | 457-470 | 106.8-107.9 |  | 12.73 | 97.52±0.82 | 105.2±0.96 |
| MT_EaHT | 10 | 191.0 | 9.1 | chr10.94.5 | chr10.95.5 | 166-198 | 9.3-9.5 | *oy1* | 15.41 | 113.6±1.04 | 100.69±1.6 |
| Ratio_EaHT | 10 | 178.0 | 11.3 | chr10.70.5 | chr10.72 | 172-197 | 6.8-6.9 |  | 18.66 | 1.12±0 | 1.01±0.01 |
| WT_Ea2Fl | 1 | 780.6 | 4.4 | chr01.1874.5 | chr01.1875.5 | 774-791 | 188.9-189.6 |  | 7.79 | 121.89±0.65 | 117.17±0.78 |
|  | 9 | 236.6 | 4.0 | chr09.130.5 | chr09.132 | 230-252 | 12.7-12.9 |  | 7.02 | 118.63±0.58 | 123.64±0.98 |
| MT_Ea2Fl | 10 | 184.0 | 12.4 | chr10.80.5 | chr10.82.5 | 179-195 | 7.9-8.1 |  | 20.43 | 124.38±0.74 | 114.35±1 |
| Ratio_Ea2Fl | 10 | 187.0 | 16.9 | chr10.88.5 | chr10.90.5 | 175-195 | 8.7-8.9 | *oy1* | 26.59 | 1.04±0 | 0.94±0 |

^a^Chromosome location of a QTL; ^b^Peak position of the QTL; ^c^LOD score at the peak of a given QTL

^d^2-LOD interval of the QTL in terms of genetic position in centiMorgan(cM); ^e^Physical position (in Mbp) of 2-LOD flanking markers from B73 RefGenV4; ^f^Phenotypic variance explained by the QTL at the peak position as estimated by regression and reported as an R^2^ value*100; ^g^Mean and standard error of the trait with B73 and Mo17 genotype at the peak marker of the detected QTL

Table S9. Morphometric analysis of mature tassels in the F_1_ progenies of the B73-like NILs and Mo17-like NIL crossed to *Oy1-N1989*/+:B73.

| Background Identifier | Genotype | TBN | BN2 | SecBN | L1 | L2 | L3 | TopBLen | MidBLen | LowBLen | AvgBLen |
| --- | --- | --- | --- | --- | --- | --- | --- | --- | --- | --- | --- |
| B73_NILs (*vey1^B73^*) | *Oy1-N1989/vey1^B73^*:B73 | 7.56±1.12bc | 1.43±0.39bc | 1.45±0.42bce | 436.3±36.13d | 225.52±23.83f | 101.77±11.89bc | 105.03±20.71d | 149.77±23.29e | 184.67±31.75d | 146.49±23.19e |
| B73_NILs (*vey1^Mo17^*) | *Oy1-N1989/vey1^Mo17^*:B73 | 3.94±0.68f | 0.3±0.36f | 0.3±0.36g | 266.63±73.88f | 132.56±41.75g | 60.18±14.29f | 68.33±26.75e | 88.38±30.98f | 99.84±33.47e | 84.19±28.64f |
| Mo17_NILs (*vey1^B73^*) | *Oy1-N1989/vey1^B73^*:Mo17/B73 | 6.86±1.83abce | 1.89±0.17abd | 1.94±0.14abcd | 538.78±28.3abce | 335.83±23.7a | 108.67±10.28abc | 167.89±26.13ab | 237.56±27.08ab | 247.22±18.84abc | 217.56±20.07ab |
| Mo17_NILs (*vey1^Mo17^*) | *Oy1-N1989/vey1^Mo17^*:Mo17/B73 | 5.24±0.89d | 0.83±0.37e | 0.84±0.38f | 439.83±57.69d | 279.96±44.92bce | 84.59±12.35de | 120.12±31.94cd | 163.31±37.49de | 178.39±33.98d | 153.94±32.4de |
| B73_NILs (*vey1^B73^*) | *vey1^B73^/vey1^B73^*:B73 | 8.38±0.93a | 1.78±0.41ad | 1.89±0.52ad | 463.39±25.61d | 243.37±16df | 110.5±8.11a | 120.64±14.05cd | 174.99±17.9cd | 218.76±22.06c | 171.46±13.84cd |
| B73_ NILs (*vey1^Mo17^*) | *vey1^Mo17^/vey1^B73^*:B73 | 7.4±0.82bc | 1.72±0.39abd | 1.81±0.42abcd | 473.17±59.97cd | 260.58±21.02cde | 108.5±10.15abc | 137.17±19.06bc | 192.32±24.7c | 233.22±30.68c | 187.57±21.03bc |
| Mo17_ NILs (*vey1^B73^*) | *vey1^Mo17^/vey1^B73^*:Mo17/B73 | 7.53±2.02abc | 1.81±0.22abcd | 1.92±0.29abcde | 545.94±48.88abe | 337.94±17.17a | 115.83±11.4ab | 191.89±16.01a | 248.89±27.42a | 278.78±17.41ab | 239.85±14.77a |
| Mo17_ NILs (*vey1^Mo17^*) | *vey1^Mo17^*/*vey1^B73^*:Mo17/B73 | 8.09±1.28ab | 2.07±0.43a | 2.14±0.49a | 562.04±28.41a | 341.64±16.13a | 117.36±15.54a | 192.38±22.82a | 248.31±27.11a | 285.69±23.17a | 242.13±21.04a |

Table S10. Tiller count in crosses of *Oy1-N1989*/+:B73 with *Tlr1* homozygotes and P39 inbred line.

| Genotype | Background | Sample size (n) | Tiller number |
| --- | --- | --- | --- |
| *+/+;Tlr1/+* | Hi27 x B73 | 14 | 1.7±0.5^a^ |
| *Oy1-N1989/+;Tlr1/+* | Hi27 x B73 | 19 | 0.1±0.3^b^ |
| *+/+* | P39 x B73 | 8 | 1.8±0.9^a^ |
| *Oy1-N1989/+* | P39 x B73 | 9 | 0±0^b^ |

Data is presented as average ± standard deviation. Connecting letter report indicates statistical significant differences between siblings within one genetic background using student’s t-test at p<0.01
